# Evidence accumulation, not “self-control,” explains dorsolateral prefrontal activation during normative choice

**DOI:** 10.1101/2020.10.06.328476

**Authors:** Cendri A. Hutcherson, Anita Tusche

## Abstract

What role do cognitive control regions like the dorsolateral prefrontal cortex (dlPFC) play in normative behavior (e.g., generosity, healthy eating)? Some models suggest that dlPFC activation during normative choice reflects the use of control to overcome default hedonistic preferences. Here, we develop an alternative account, showing that an *attribute-based neural drift diffusion model (anDDM)* predicts trial-by-trial variation in dlPFC response across three fMRI studies and two self-control contexts (altruistic sacrifice and healthy eating). Using the anDDM to simulate a variety of self-control dilemmas generated a novel prediction: although dlPFC activity might *typically* increase for norm-consistent choices, deliberate self-regulation focused on normative goals should *decrease* or even *reverse* this pattern (i.e., greater dlPFC response for hedonic, self-interested choices). We confirmed these predictions in both altruistic and dietary choice contexts. Our results suggest that dlPFC response during normative choice may depend more on value-based evidence accumulation than inhibition of our baser instincts.

## Introduction

Self-control dilemmas typically involve trade offs between short-term, hedonic considerations and longer-term or more abstract standards and values. For example, social interactions often force an individual to weigh self-interest against norms favoring equity and other-regard. Similarly, dietary decisions often require weighing the immediate pleasure of consumption against personal standards or societal norms favoring healthy eating. Understanding when, why, and how people choose normatively-preferred responses (e.g., generosity over selfishness, healthy over unhealthy eating, etc.) has represented a central goal of the decision sciences for decades. What neural and computational processes must be engaged to support more normative behavior? What makes such choices frequently feel so conflicted and effortful, and how can we make them easier? To what extent does following social or personal norms depend on activation in brain regions associated with cognitive control, such as the dorsolateral prefrontal cortex (dlPFC)?

Previous research has provided a wealth of evidence suggesting that the dlPFC may promote normative choices in both the social and non-social domain. For instance, compared to unhealthy food choices, healthier choices in successful dieters were accompanied by greater activation in a posterior region of the dlPFC^1^. Greater dlPFC response in a similar region has also been observed when individuals make normatively-favored choices in both social decision making^2, 3^ and intertemporal choice^4, 5^. Moreover, activation in the dlPFC increases when individuals explicitly focus on eating healthy^6^ or on decreasing craving for food^7^. Electrical disruption of this area also decreases patience^8^ and reduces normative behavior in social contexts like the Ultimatum game^9^. Collectively, these results support the notion that the dlPFC may be recruited to modulate values or bias choices in favor of normative responses, perhaps especially when those responses conflict with default preferences.

Yet a variety of results seem inconsistent with this view. For example, researchers often fail to observe increased dlPFC recruitment when individuals make pro-social or intertemporally normative choices^10–12^. Moreover, electrical disruption of the dlPFC has been observed both to *decrease* appetitive valuation of foods^13^, and *increase* generous behavior in the Dictator Game^9^. Such findings conflict with the idea that this region consistently promotes normative concerns over immediate, hedonistic desires. Thus, how to predict whether and when one might observe a positive association between dlPFC response and choices typically associated with successful self-control remains unclear.

Here, we propose a computational account of fMRI BOLD response in the dlPFC that may resolve many of these apparent inconsistencies. This account draws on prior research in both perceptual and value-based decision making, which consistently finds that the posterior dlPFC region associated with normative “self-control success” also activates during choices that are more difficult to discriminate in simple perceptual and value-based choices lacking a self-control conflict, e.g., ^14–16^. Our account is also inspired by findings that the dlPFC may be one hub in a larger neural circuit (encompassing additional regions like the dorsal anterior cingulate cortex [dACC], supplementary motor area [SMA] and inferior frontal gyrus/anterior insula [IFG/aIns]) that selects actions for execution using a process of evidence accumulation and lateral inhibition among competing action representations^17, 18^. Based on this evidence, we developed a computational model of self-control dilemmas that successfully predicts not only when an individual will choose in normative rather than hedonistic fashion, but also when, why, and to what degree response in the dlPFC will be recruited during that process. We note also that, although we focus here on the dlPFC, our model also applies in theory when observing similar relationships to other brain areas frequently associated with conflict and cognitive control, including regions of the IFG/aIns and dACC.

As with similar models of simple perceptual and value-based choices, our *attribute-based neural drift diffusion model* (anDDM) assumes that the brain makes decisions through a process of value-based attribute integration and competition (Figure 1). More specifically, choices are resolved via competitive interactions between neuronal populations that output responses based on accumulated information about the value of choice attributes, weighted by their momentary goal relevance. Some of these attributes are associated with hedonism (e.g., self-regarding concerns in altruistic choice) and some are associated with social norms and standards for behavior (e.g. other-regarding concerns). For expository purposes, we refer to these respectively as hedonic and normative attributes. Intuitively, whether our computational algorithm makes a hedonistic or normative choice depends not only on the magnitude of hedonic and normative attributes, but also on their weight: higher weights on normative attributes lead to more norm-consistent responses.

**Figure 1.**
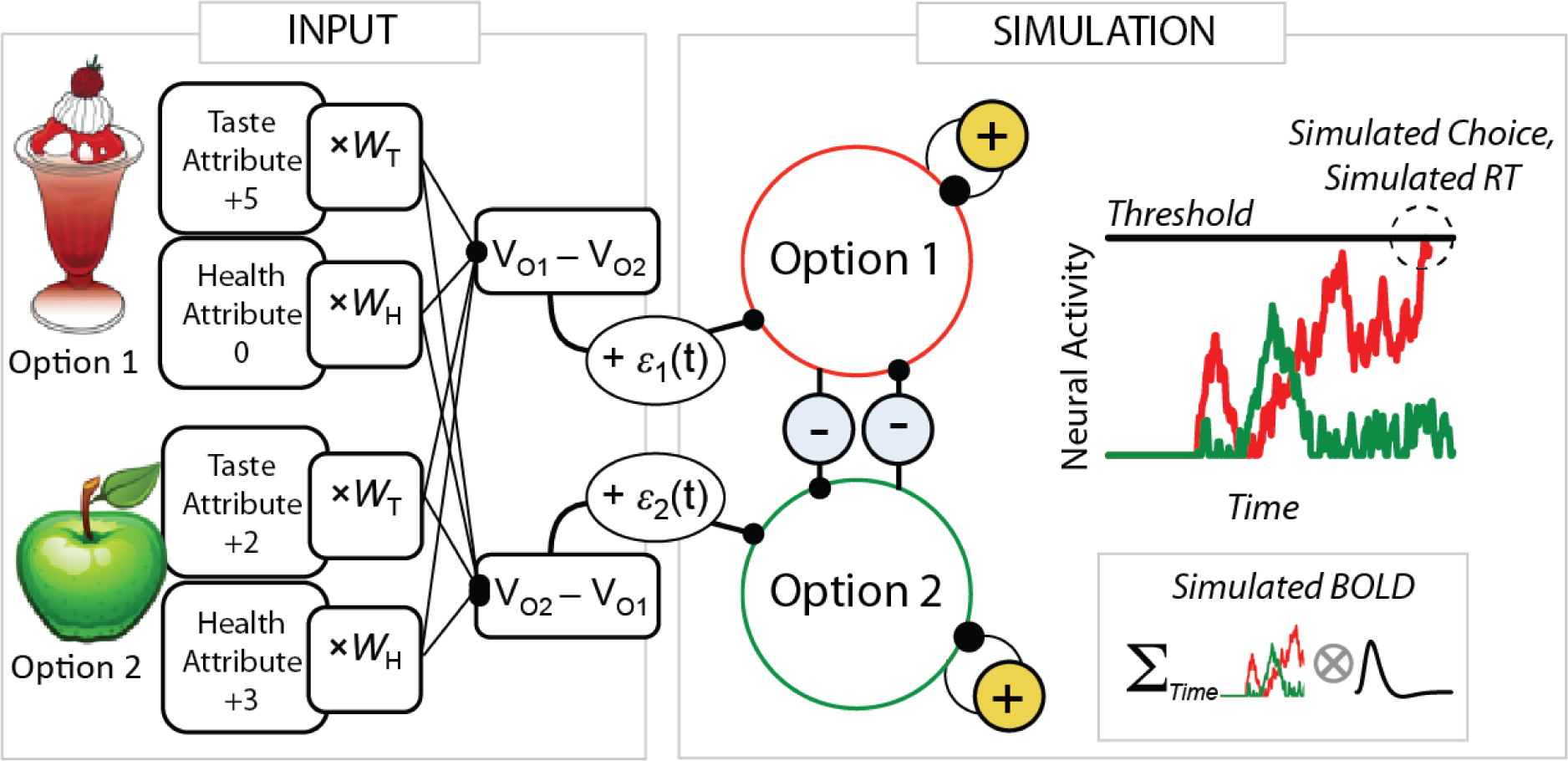
Attribute-based neural drift diffusion model (anDDM) of normative choice. Each option’s hedonic and normative attributes (e.g., tastiness = +5 and healthiness = 0 for the sundae) are weighted by their current importance (e.g., w_Taste_ [w_T_] and w_Health_ [w_H_]) and summed to construct relative option values [V_O1_ – V_O2_]. These values, corrupted by momentary noise at time t [ε1(t)], serve as the external inputs to two mutually inhibitory neuronal pools representing the two options. Neural activation in these two pools (red and green lines in upper right plot) accumulates over time until one hits a predefined threshold, determining both the simulated response time (RT) and the simulated choice. Choices are classified as normative if the option with higher normative attribute value (in this case, higher healthiness, i.e. the apple in option 2) is selected. The sum of neural activation across the two pools can be used to simulate expected neural signals at the time of choice, and can be convolved with the canonical hemodynamic response function to construct a predicted BOLD signal for each choice (lower right inset).

What role does the dlPFC play in the anDDM? The observation of increased posterior dlPFC response when people choose consistently with normatively favored goals (e.g., healthy over unhealthy choices) has been taken to suggest that this region acts either to modulate the processing of attribute values or their weights in favor of normatively-favored goals^1, 6^, or to inhibit hedonistic reward-related responding^19, 20^. In contrast, we propose that activity in this region reflects processes related to the *response selection stage* of decisions. This suggests that dlPFC response during normative choice represents a downstream consequence of valuation processes, rather than a direct causal influence upon them. To support this argument, we use the anDDM to simulate when and why we might observe greater activity in the dlPFC (and regions with similar response profiles) when resolving a choice. As we describe below, these simulations suggest that normative choices should be associated with greater neural activation in the dlPFC only when two things are true: hedonic attribute values *directly oppose* normative attribute values, and hedonic attributes receive *more weight* as inputs to the anDDM. In contrast, when normative attributes receive more weight, *hedonistic* choices should produce greater activity in the dlPFC and other areas associated with response selection.

We then used these observations to make two predictions. First, if people by default favor hedonic over normative attributes, then most studies will observe greater dlPFC response when people choose the normatively-favored option. This prediction does not strongly distinguish our account from alternatives. However, our model makes a second, more novel prediction: if a normally hedonistic decision maker focuses on normative goals, this should *reduce* activation in the dlPFC when choosing the normatively-favored option. A straightforward reading of an attribute-weighting account predicts the opposite: a normally hedonistic individual who deliberately attempts to focus on normative responding should show *increased* activation in the dlPFC in order to alter attribute weighting in favor of normative goals^19, 21^. We test these two alternative predictions across three studies and two canonical self-control contexts in which people frequently struggle to align their actual behaviors with normative goals: altruistic and dietary choice. In all cases, results strongly supported the predictions of the anDDM. These findings raise new and important questions regarding the role of the dlPFC– and effortful self-control more generally – in promoting normative choice.

## Results

### Simulating the dilemma of self-control

Although self-control dilemmas can take a variety of forms, for expository purposes we here take a single, typical self-control dilemma: a decision maker deciding whether to indulge in a decadent snack or opt for something healthier. This example allows us to capture two critical features: first, self-control dilemmas typically involve making decisions about options that vary in the magnitude or value of hedonic and normative attributes (e.g. tastiness and healthiness). Second, the decision-maker must weigh these attributes based on goals that can vary in their relative strength at different times. At a nice restaurant, tastiness may be prioritized. When trying to lose weight, healthiness is prioritized. We used simulations to explicitly capture these two features.

Simulations were realized using a neural network instantiation of our anDDM^18^ where choices result from dynamic interactions between two separate but intermingled pools of neurons representing the different options under consideration (Figure 1). Activation in each pool accumulates noisily based on a combination of external inputs from hedonic and normative attributes weighted by their current subjective importance, inhibitory inputs from the other pool, and recurrent self-stimulation (see Methods for details). This model generated predictions for how *magnitudes* and *weights* for hedonic and normative attributes influence the likelihood of a virtuous (i.e., healthy) choice, response time [RT], and neural response. These simulations yielded three key observations about behavior and neural response, which we describe in the context of food choice but apply in theory across any self-control dilemma that requires weighing hedonic rewards against normative values and goals.

#### Observation 1

*The likelihood of a normative choice depends on the value of hedonic and normative attributes.* To capture the idea that some choices (e.g. ice cream vs. Brussels sprouts) represent more of a self-control conflict than others (e.g. strawberries vs. lard), we simulated a single decision maker facing choices between hypothetical options that independently varied the relative value of normative and hedonic attributes (e.g. the foods’ relative healthiness and tastiness). In the context of food choice, we classified a simulated choice as normative (healthy) when the simulation selected the option with higher healthiness. Choices were classified as hedonistic (unhealthy) otherwise. To determine the effect of current behavioral goals, we simulated the decision maker’s choices for a variety of different weights on healthiness (*w_Health_*) and tastiness (*w_Taste_*).

Figure 2a illustrates how variation in tastiness and healthiness of an option relative to the alternative affects a decision maker’s *general* propensity to make a healthy choice (i.e., averaging over different instances of *w_Taste_* and *w_Health_*). As can be seen, the magnitude and sign of the two attributes matters: she tends to choose more healthily when one option dominates on both healthiness and tastiness (no-conflict trials). She chooses less healthily when one option is tastier while the other is healthier (conflict trials). She is least likely to choose normatively when the difference in tastiness is large and the difference in healthiness is small. Thus, our simulations make the commonsense prediction that attribute values matter in determining the overall likelihood that an individual makes a healthy/normative choice.

**Figure 2.**
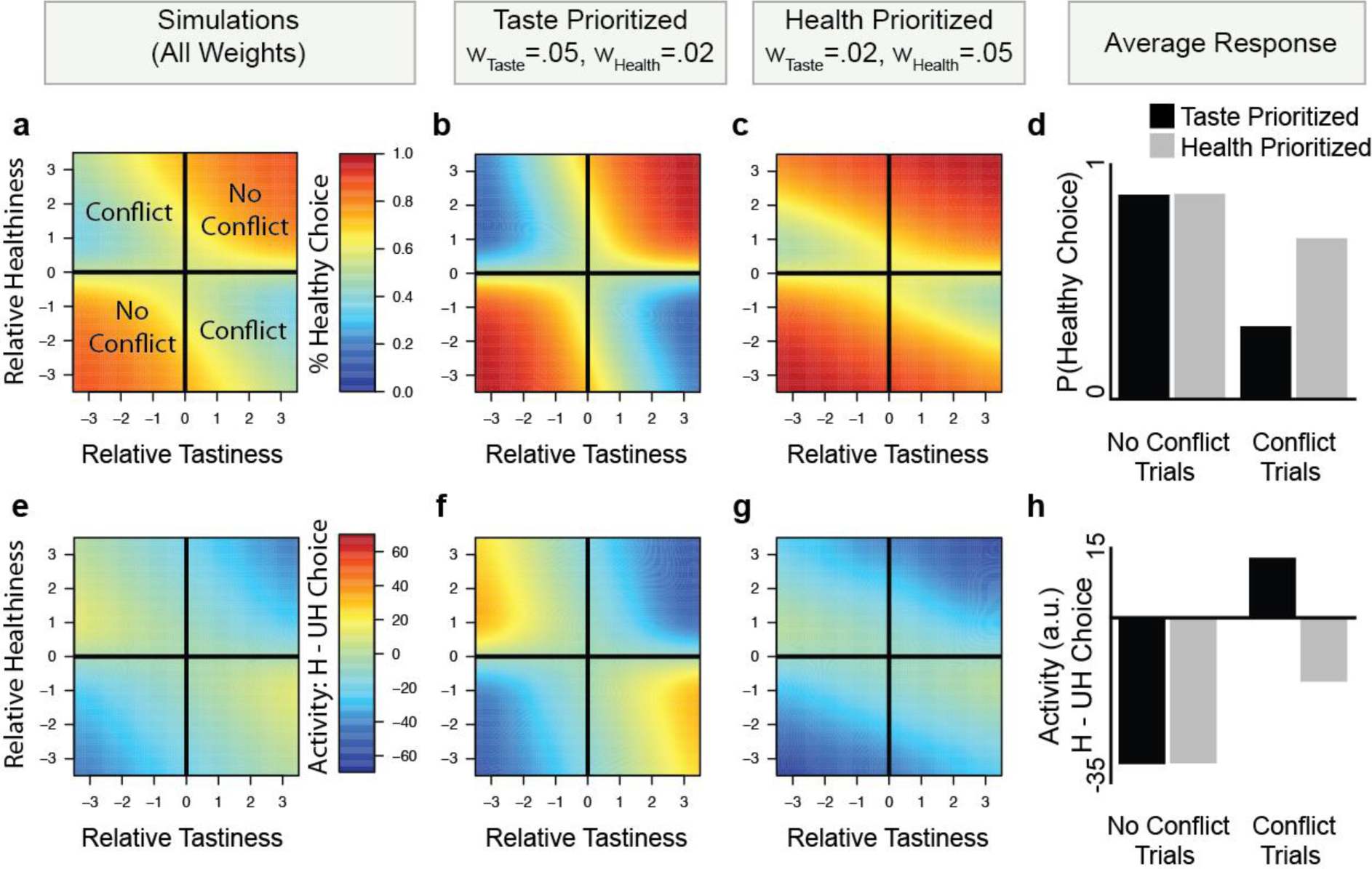
Simulating the dilemma of self-control. Top: The computational model can be used to simulate decision making for any self-control context requiring an integration of normative and hedonistic considerations (healthy eating displayed). (**a**) On average across multiple different goals, the likelihood of a healthy choice depends on the relative attribute values of one option vs. another, and is less likely when tastiness and healthiness conflict. Warmer colors indicate a higher likelihood of choosing the healthier option. Specific goals (**b**) prioritizing tastiness or (**c**) prioritizing healthiness alter the overall frequency of healthy choice, although in both contexts unhealthy choices are more likely for large differences in tastiness and small differences in healthiness. (**d**) The overall likelihood of a healthy choice (averaged for all combinations of conflict or no conflict choices). Goals prioritizing tastiness (black bars) produce fewer healthy choices than goals prioritizing healthiness (gray bars), but only when tastiness and healthiness conflict. Bottom: **e**-**g**) The computational model can also simulate expected neural activity (i.e. aggregate activity in the two neuronal pools, summed over decision time: *∑_Time_ Option1 + Option2*) when choosing healthy [H] or unhealthy [UH] options, as a function of relative option values and different goals. Warmer colors indicate more activity when a healthy choice was made (i.e., Activity _H_ > Activity _UH_). **h**) Overall difference in neural activity for H compared to UH choices for goals prioritizing tastiness (black bars) and healthiness (gray bars), divided as a function of attribute conflict. In no conflict trials, healthy choices elicit less activity regardless of goal (i.e. Activity _H_ < Activity _UH_). In conflict trials, however, healthy choices elicit more activity (i.e. Activity _H_ > Activity _UH_), but only when goals prioritize tastiness. Identical results are obtained when substituting RT for neural response (see Supplementary Figure 1).

#### Observation 2

*The likelihood of a normative choice depends on weights given to normative and hedonic attributes.* We next attempted to capture the idea that an individual might vary from context to context in the goals that they prioritize, and that the essence of self-control is to prioritize (i.e., assign a higher weight to) normative attributes like healthiness, or to deprioritize (i.e., assign a lower weight to) hedonic attributes like tastiness. We thus simulated the decision maker in different goal states by assuming different weights on hedonic and normative attributes (i.e. tastiness and healthiness). We show two example simulations in Figure 2b-d. Unsurprisingly, the decision maker chooses healthily less frequently when weight on tastiness is higher than weight on healthiness. However, these differences are starkest in conflict trials, and essentially vanish for no-conflict trials (Figure 1d).

#### Observation 3

*Normative choices result in higher neural response only if attributes conflict and the decision maker weights hedonic attributes more.* The last and most important goal of our computational model simulations was to examine how neural response in a cognitive control region like the dlPFC (assuming its activity correlates with the anDDM) might depend on weights given to hedonic and normative attributes (Figure 2e-h). We characterized this simulated response as aggregate activity of the two neuronal pools, summed over the duration of the choice, as this is what would contribute to observable BOLD responses.

Comparing differences in simulated neural response for healthy and unhealthy choices yields two important conclusions. First, when options do not conflict on healthiness or tastiness (i.e. one option is better on both), healthy choices generally elicit *less* activity than unhealthy ones (Figure 2e). Notably, for no-conflict trials this holds true irrespective of whether a decision maker is currently prioritizing tastiness or healthiness (Figure 2f-g). Second, and more importantly, when attributes *conflict*, network activity during healthy vs. unhealthy choices shows a striking dependence on an individual’s goals (i.e. the relative balance of *w_Health_* and *w_Taste_*). In conflict trials, hedonism-favoring goals (i.e., w_Taste_ > w_Health_) result in higher activity on average when choosing healthily (Figure 2h). This difference becomes exaggerated as the magnitudes of tastiness and healthiness increase (Figure 2f). In contrast, when goals prioritize normative attributes like healthiness (i.e., w_Health_ > w_Taste_), simulated neural responses are *lower* on average for healthy compared to unhealthy choices (Figure 2g,h). Thus, neural response is positively associated with normative choice (i.e., greater neural activity to choose normatively instead of hedonistically) only when the decision maker places a higher weight on hedonistic than normative attributes. The same is true of simulated RTs, which are often used as a proxy for both choice difficulty and the presence of control (Supplementary Figure 1). Thus, in the anDDM the observation that normative choices activate brain areas associated with cognitive control might simply indicate that hedonic attributes are currently weighted more highly.

### Testing computational predictions using fMRI data

#### The anDDM accurately predicts dlPFC activity across a variety of contexts

It is currently unknown whether activity in the dlPFC region frequently associated with self-control might reflect activation patterns in the anDDM in the same manner as simple choice^18^. We thus began by verifying that trial-by-trial simulated neural activity in the anDDM correlated with activity in this region for complex, multi-attribute choices typical of different real-world self-control dilemmas. Note that, while this correlation could occur because the dlPFC performs the precise computations carried out by the anDDM, such a correlation could also occur if the dlPFC performs separate computational functions that activate proportionally to anDDM activity. In either case, we would expect trial-by-trial activity of the dlPFC to correlate with predictions of the anDDM.

Our analysis focused on three previously-collected fMRI datasets^22, 23^ (see Methods for details). Study 1 (N = 51) and Study 2 (N = 49) utilized an Altruistic Choice Task trading off different monetary outcomes for self and an anonymous partner in a modified version of a Dictator game (Figure 3a, b, see Methods for details). Study 3, completed on a subset of participants from Study 2 (N = 36), utilized a Food Choice Task (Figure 3c) with different foods varying in tastiness and healthiness. In Study 1, choices were made with the instruction to simply choose the most-preferred option. In Studies 2 and 3, participants made choices in three separate conditions that manipulated goals/attribute weights by instructing participants to focus on different normative or hedonistic attributes (a point we return to below). Studies 1 and 2 involved only trials involving conflict between hedonic and normative attributes. Study 3 included trials both with and without such conflict.

**Figure 3.**
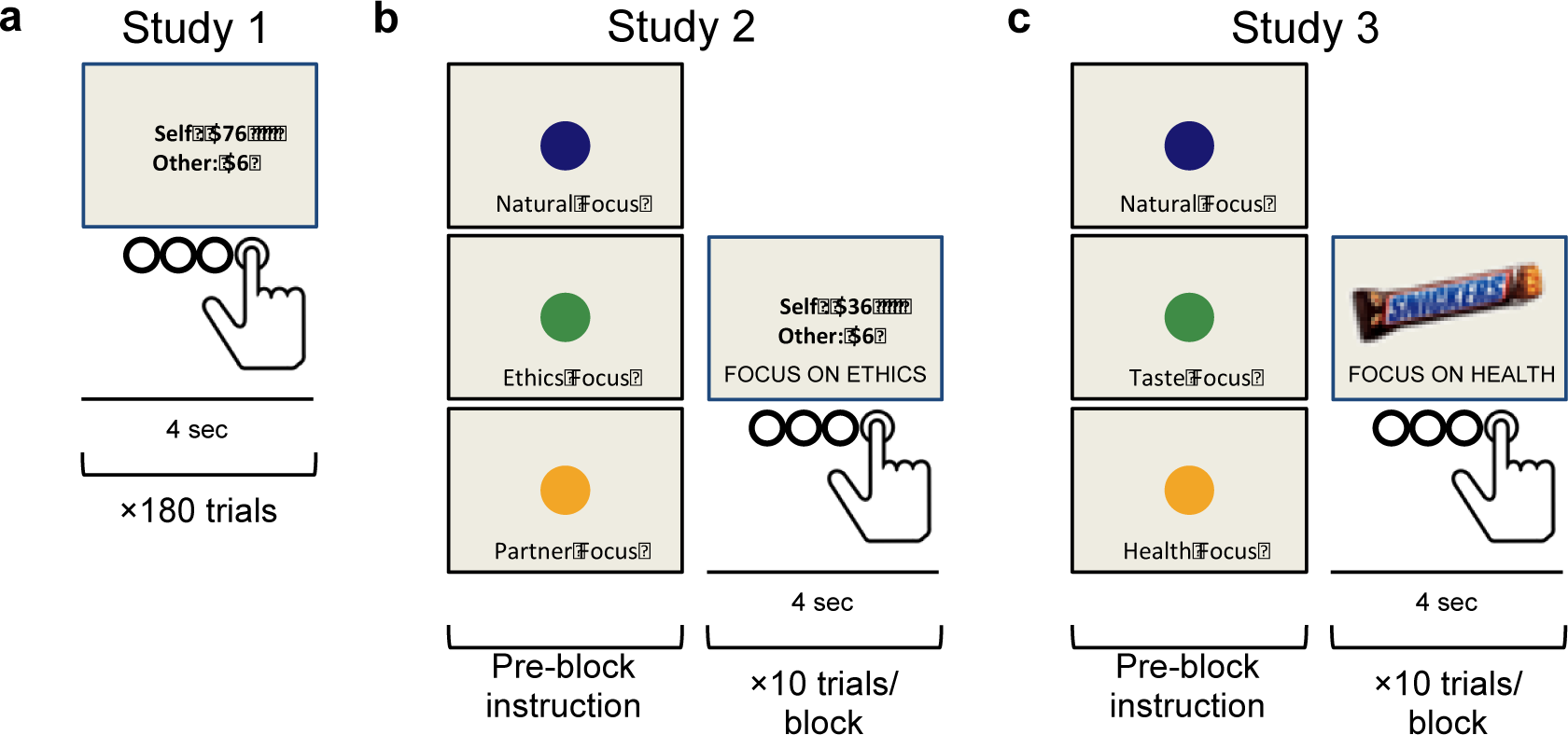
FMRI task designs. **(a)** In Study 1, participants made choices involving tradeoffs between monetary payoff for another person ($Other; normative attribute) and for themselves ($Self; hedonic attribute) in an Altruistic Choice Task. **(b)** In Study 2, participants made choices similar to the Altruistic Choice Task in Study 1, while we manipulated the *weights* on normative and hedonic attributes using instructions presented at the beginning of each task block. These instructions asked participants to focus on different pro-social motivations (ethical considerations, partner’s feelings) as they made their choice. **(c)** In Study 3, we examined the generalizability of the model-based predictions in another choice domain. Here, we manipulated weights on food’s healthiness (normative attribute) and tastiness (hedonic attribute) using a Food Choice Task. In all studies, participants had 4 seconds to decide, and gave their response on a 4-point scale from “Strong No” to “Strong Yes”.

We predicted that dlPFC activity should correlate parametrically with simulated activity of the anDDM during self-control dilemmas. To test this notion, we first fit computational parameters of the anDDM to each participant’s behavior (see Supplemental Figure 2 for model fits). We then asked whether parametric variation in the measured BOLD signals within the dlPFC ROI correlated with simulated response across all three fMRI studies (see Methods for detail). To this end, data of each study were thresholded at a voxel-wise *P* < .001, and a cluster-defining threshold of *P* < .05, small-volume corrected within a 10-mm spherical region of interest (ROI) centered on the peak coordinates of activity for the contrast of normative (healthy) vs. hedonistic (unhealthy) choice in a previous study of self-control in dieters^1^. The results of a three-way conjunction at this a priori threshold show that anDDM responses correlate with activation in the dlPFC across all three data sets (Figure 4a, center-of-mass x = −56, y = 19, z = 21). Results for our key questions reported below (Figure 4 e-f) are based on the dlPFC cluster identified in this conjunction analysis. Supplemental analyses confirmed that simulated activity of the anDDM covaried with observed BOLD responses in the DLPFC in each condition of Study 2 and 3.

**Figure 4.**
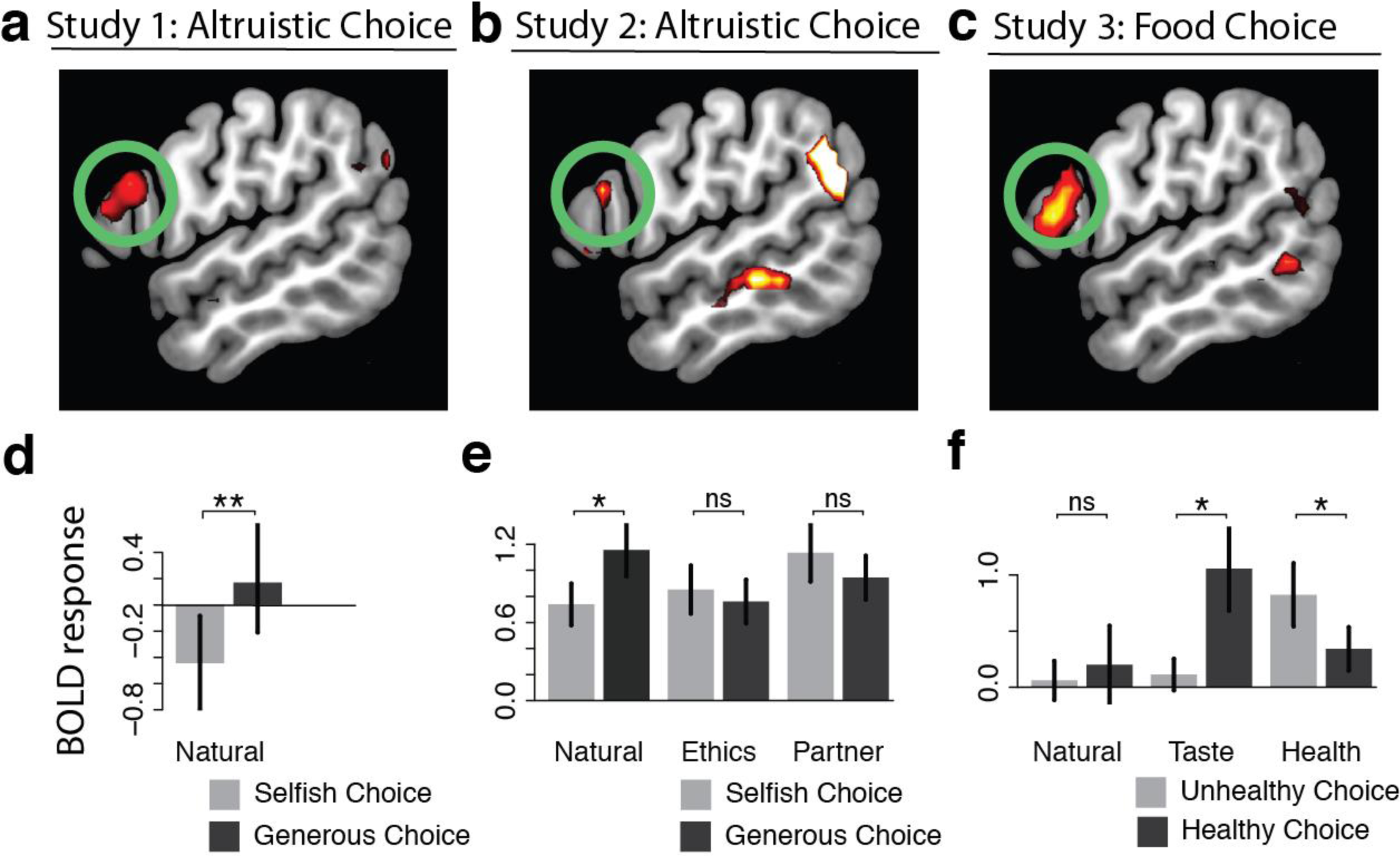
BOLD responses in the left dlPFC during self-control dilemmas. Top: Trial-by-trial BOLD response in the dlPFC correlates with predicted activity of the anDDM across three separate studies, including during both altruistic choice **(a, b)** and during dietary choice **(c)**. All maps thresholded at *P* < .001 uncorrected for display purposes. Bottom: Within the dlPFC ROI defined by the three-way conjunction of anDDM response across all studies, BOLD response during normative choice (black) vs. hedonistic choice (light gray) when attributes conflict, in **d**) Study 1 for all trials, as well as in **e**) Study 2 and **f**) Study 3 as a function of regulatory goals. As predicted, normative choices activate the dlPFC, but only when goals result in a greater weight on hedonistic than normative attributes. * *P* < .05; ** *P <* .01.

Intriguingly, although they are not the focus of this study, we also observed a whole-brain corrected conjunction of activation across all three studies in two other regions often associated with conflict and cognitive control: the dorsal anterior cingulate cortex (dACC) and anterior insula/inferior frontal gyrus (*P*s < .001, whole-brain corrected across all three studies, Supplemental Figures S3 and S4). No other regions showed a similarly consistent, three-way conjunction across all three studies.

### Recruitment of the dlPFC when choosing normatively only occurs when goals are hedonistic and attributes conflict (Observation #3)

The preceding analysis confirmed that activity in the left dlPFC covaries with predicted activity simulated in the anDDM in three independent fMRI studies. We next confirmed the central prediction of our simulations concerning the relationship between normative choices and activity in the dlPFC. In particular, models suggesting that the dlPFC promotes normative choices^1, 6, 20^ imply that norm-consistent choices should be accompanied by greater activation in the dlPFC (as has been observed previously). Moreover, this should be especially true when people focus on normative goals^6, 7^, since those goals support norm-sensitive behavior and might require the override of default hedonistic preferences^19, 24^. The anDDM makes the opposite prediction. While neural activity in the model (and by extension the dlPFC) *can* be higher for normative compared to hedonistic choices, this should be true only when goals lead to stronger weighting of hedonic attributes and attribute values conflict (c.f. Figure 2h). Thus, if a regulatory focus on normative attributes increases their weight in the evidence accumulation process, this should increase normative choices, but result in *lower*, not higher, neural activity for those choices. We tested these predictions by performing a region-of-interest (ROI) analysis in the dlPFC region identified by the three-way conjunction above, examining the contrast of activity for normative compared to hedonistic choices in different contexts. In Study 1 (altruistic choice) this involved choices made only during natural, unregulated decision making. In Study 2 (altruistic choice) and Study 3 (food choice) we examined choices made under different regulatory goals that were designed to increase or decrease weights on hedonic and normative attributes (i.e. self and other in altruistic choice, tastiness and healthiness in food choice).

#### Generous vs. selfish choices (Study 1)

In Study 1, choices were defined as normative (i.e., generous) if the participant selected the option with less money for themselves and more money for their partner. Choices were defined as hedonistic (i.e., selfish) otherwise. Weights from the best-fitting model parameters indicated that subjects naturally placed more weight on their own outcomes (mean *w_Self_* = .0036±.0011s.d.) than the other person’s outcomes (mean *w_Other_ =* .0008±.0015, paired-t_50_ = 12.37, P = 2.2×10^-16^) or on fairness (i.e., |Self – Other|, mean *w_Fairness_ =* .0008±.001, paired-t_50_ = 8.30, P = 7.82×10^-11^). Given the higher weight on self-interest, a hedonic attribute, and the fact that all trials in this study involved conflict between normative and hedonic attributes, we predicted that we should observe greater neural response when people chose generously. An ROI analysis of BOLD response in the dlPFC for generous vs. selfish choices strongly supported this prediction (Figure 4d, paired-t_43_ = 2.98, *P* = .005). A whole-brain analysis confirmed that this pattern was specific to the dlPFC, as well as the dACC and insula/IFG regions also associated with the anDDM, rather than a general property of neural activity (see Supplementary Table 3 for details).

#### Regulatory effects on generous vs. selfish responding (Study 2)

In Study 2 (also anonymous altruistic decision making and conflict trials only), we sought to replicate and extend these results. More specifically, we sought to test the anDDM prediction that if regulatory goals increase the weight on normative attributes, this should result in *decreased* activation in the dlPFC when choosing normatively. To manipulate weights on hedonic and normative attributes, we used an instructed cognitive regulation manipulation in which we asked participants on different trials either to “Respond Naturally” (mirroring the natural preferences expressed by participants in Study 1) or to focus on one of two different goals (“Focus on Ethics” [Ethics], “Focus on your Partner’s Feelings” [Partner]) that both emphasize normative attributes, but in different ways (see Methods for details). To confirm that the manipulation influenced attribute weights, we performed one-way repeated-measures ANOVAs with condition (Natural, Ethics, Partner) as a fixed effect and best-fitting attribute weight parameters wSelf, wOther, and wFairness as dependent variables. This analysis confirmed that our manipulation yielded significantly different weights on the attributes across the conditions (all F_2,96_ > 13.54, all P < 6.59×10^-6^, see Methods for details of model fitting). As expected, weights for self-interest (a hedonic attribute, w_Self_) were highest in the Natural condition (M_Natural_ = .0073±.0035 s.d.), lower in the Ethics condition (MEthics = .0061±.0047), and lowest in the Partner condition (M_Partner_ = .0037±.0065). By contrast, weights on the partner’s outcomes and fairness (attributes related more strongly to social norms) increased with regulation (wOther: M_Natural_ = .0010±.0038, M_Ethics_ = .0041±.0045, M_Partner_ = .0051±.0038; wFairness: M_Natural_ = .0017±.0033, M_Ethics_ = .0053±.0046, M_Partner_ = .0024±.0035).

Having confirmed that the regulatory focus manipulation altered weights on hedonic and normative attributes, we next asked if this manipulation affected BOLD response during generous vs. selfish choice in the dlPFC, consistent with predictions of the anDDM. In particular, given that all trials involved conflict between normative and hedonic attributes, we predicted that in the Natural condition, where participants generally placed higher weight on self-interest (a hedonic attribute), *generous* choices should elicit higher activation. In contrast, in the Partner condition, which elicited higher weight on normative attributes (i.e., other’s outcomes and fairness), *selfish* choices should elicit the greatest activity in the dlPFC. The Ethics condition, which elicited similar weights across the attributes, should lie in between.

To test these predictions, we performed one-way repeated measures ANOVAs with condition (Natural, Ethics, Partner) as a fixed effect and average BOLD response in the dlPFC ROI for the contrast of generous vs. selfish choice as the dependent variable. This analysis revealed a significant effect of condition on dlPFC response (F_2,96_ = 4.67, P = .01). Post-hoc planned comparisons confirmed that in the Natural condition, generous choices elicited significantly greater activity in the dlPFC (P = .04, Figure 4e), replicating the observed difference during Natural choices of Study 1. By contrast, in the Ethics and Partner focus conditions, generous choices no longer elicited significantly greater activation. Instead, *selfish* choices elicited *greater* activation, although the effect did not reach statistical significance. Thus, in the same individuals, the association between generous choices and *higher* activation in the dlPFC depended on whether goals emphasized selfishness rather than social norms (Figure 4e). Supplemental whole-brain analyses confirmed these findings (see Supplementary Results, and Supplementary Table 3 for details).

#### Regulatory effects on healthy vs. unhealthy choice (Study 3)

In Study 3, we sought to replicate the finding that a regulatory focus on normative attributes reduces activation in the dlPFC, but in a new, non-social domain: healthy eating. During the Food Choice Task in Study 3, we manipulated attribute weights by instructing participants either to “Respond Naturally”, “Focus on Health”, or “Focus on Taste” while making their choice. Normative (i.e., healthy) choices were defined as selecting the food with higher subjectively perceived healthiness (see Methods for details). Note that the “Focus on Health” instruction aimed to increase weight on healthiness (w_Health_), a normative attribute. Extending results of Study 2, the “Focus on Taste” condition was designed to enhance the weight on tastiness (w_Taste_), the hedonic attribute, which should preserve or even enhance the difficulty of normative choices that we observed in natural choice settings in study 1 and 2. This allowed us to verify that our findings are specifically driven by changes in weights, not simply because we asked participants to perform a cognitive task.

To confirm that the regulatory manipulation influenced attribute weights, we performed one-way repeated-measures ANOVAs, similar to Study 2, with condition (Natural, Taste, Health) as a fixed effect and estimated attribute weight parameters w_Taste_ and w_Health_ as dependent variables. This analysis confirmed that our manipulation yielded significantly different weights on the different attributes across the conditions (all Fs > 104.2, all P < 2.2×10^-6^). As expected, weights on tastiness (a hedonic attribute) were highest in the Taste condition (MTaste = 0.0077±.0029), similar but slightly lower in the Natural condition (M_Natural_ = 0.0074±.0027) and lowest in the Health condition (MHealth = 0.002±0.0028). Weights on healthiness (a normative attribute) showed the opposite pattern, being lowest in the Taste condition (MTaste = −0.0008±0.0018), similar though slightly higher in the Natural condition (M_Natural_ = −0.0002±0.0018) and highest in the Health condition (MHealth = 0.0055±0.0034).

Given these weights, we predicted that on the subset of trials involving conflict between healthiness and tastiness, healthy compared to unhealthy choices should elicit the greatest activation in the dlPFC in the Taste condition. In contrast, *unhealthy* choices should elicit greater activation in the Health condition. The Natural condition should lie in between these two extremes, being more similar to the Taste condition. To test these predictions, we performed a one-way repeated measures ANOVA, similar to Study 2, with condition (Natural, Taste, Health) as a fixed effect and the average dlPFC BOLD response in the contrast of healthy vs. unhealthy choice (limited to trials with attribute conflict) as the dependent variable. As hypothesized, this analysis revealed a significant effect of condition on response (F = 4.269, *P* = .018). Follow-up t-tests confirmed the predicted direction of activation (Figure 4f). BOLD response during healthy compared to unhealthy choices was significantly greater in the Taste condition for the dlPFC (paired-t32 = 2.67, *P* = .01). In the Health condition by contrast, activity was significantly greater for *unhealthy* choices in the left dlPFC (paired-t34 = 2.061, *P* = .05). Response for healthy vs. unhealthy choice in the Natural condition lay in between these two extremes. Thus, in the same individuals, healthy choices could be accompanied by *higher* activation in brain regions typically associated with cognitive control (when goals emphasized hedonism), or *lower* activation (when goals emphasized health norms). Supplemental whole-brain analyses confirmed that this pattern of results was specific to the dlPFC and other regions associated with the anDDM (see Supplementary Results, and Supplementary Table 3 for details).

#### Regulatory effects in the absence of conflict (Study 3)

Our analyses so far focused on conflict trials, since simulations suggest that these trials show the biggest differences as a function of attribute weights (Figure 2). The design of Study 3, which included a subset of trials with no attribute conflict, also allowed us to test one further prediction of the anDDM. In Observation #3, we found that normative choices should only be associated with increased neural activity *when hedonic and normative attributes conflict* (Figure 2h). When attributes do *not* conflict, the anDDM predicts that normative choices should on average result in *lower* neural response. Moreover, the anDDM suggests smaller differences in response across goal contexts favoring hedonism or health norms. This suggests that, in contrast to conflicted choices, there should be less effect of regulatory focus on dlPFC response during no-conflict choices.

To test this prediction, we first performed a one-way repeated measures ANOVA with condition (Natural, Taste, Health) as a fixed effect and the average BOLD in the dlPFC for the contrast of healthy vs. unhealthy choice as the dependent variable, focusing only on the subset of trials with no conflict between tastiness and healthiness of a food (i.e., when the value of the option was positive or negative for both). As predicted, there was no significant influence of regulatory condition on the difference in neural activity between healthy and unhealthy choice (F2,68 = 0.477, *P* = .62). Given this lack of effect across conditions, we averaged the three conditions together to analyze the main effect of healthy vs. unhealthy choice. This analysis indicated that healthy choices were accompanied by non-significantly *lower* response in this region (paired-t35 = 1.51, p = .07, one-tailed). Results in other regions correlating with the anDDM, including the dACC and insula/IFG showed an even stronger pattern (see Supplemental Results for more details). In other words, as expected from model simulations, activation in the dlPFC for normative choices when normative and hedonic attributes did not conflict is generally low, and shows little to no effect of regulatory focus or the relative weight on tastiness and healthiness.

#### Regulation-related differences in overall activation (Studies 2 & 3)

Our analyses so far confirm predicted patterns of response in the dlPFC during normative choice, suggesting that altering weights on normative vs. hedonic attributes alters the association between the dlPFC and normative choice. This raises the obvious question: which regions of the brain produce these changes in weight? Some models attribute this role to the dlPFC itself, arguing that increases in activation in this area when focused on specific attributes (e.g. focusing on healthy eating) reflect computations necessary to redirect attention and alter weights. We thus interrogated the dlPFC for evidence that activation in this area during either Study 2 or Study 3 might increase generally when people focus on regulating their attention, as might be expected if this region implements changes in weights. However, we observed no effect of regulatory focus on overall response in this region in either Study 2 (F2.96 = 1.12, *P* = .33) or Study 3 (F2.70 = 1.294, *P* = .28). Thus, we found no evidence that this region activates to *drive* changes in weights.

## Discussion

When and why do normative choices (i.e., those choices that conform to abstract standards and social rules) recruit regions associated with cognitive control like the dorsolateral prefrontal cortex (dlPFC)? Simulated activity from an attribute-based neural drift diffusion model (anDDM) suggests a straightforward answer: normative behavior may only trigger the dlPFC when normative attributes conflict with hedonic ones, and the decision maker values hedonic attributes more. Across three separate fMRI studies and two different choice domains (generosity and healthy eating), we show several results that confirm predictions of the anDDM. First, we show that activation in the dlPFC correlates consistently with predicted activity of the anDDM across all contexts examined. Second, we show that even in individuals who show a natural bias towards selfishness, regulatory instructions to focus on socially normative attributes increase generosity but *reduce* dlPFC response when choosing generously. Third, this pattern replicated in the domain of healthy eating, suggesting a general principle that may apply across a variety of self-control dilemmas. Finally, we found little evidence that overall activation in the dlPFC predicted regulation-induced changes in weight. Our results provide empirical support for recent theories positing that successful self-control—defined as choosing long-term or abstract benefits over hedonic, immediate gratification^25^—depends importantly on value computations. They stand in contrast to the predictions of models of posterior dlPFC function suggesting that the strength with which the dlPFC activates during choice determines whether prepotent hedonistic responses are resisted^19, 21, 24, 26^. Our results point to a modified conceptualization of the role played by the dlPFC in promoting normative choice.

A large literature, generally consistent with models that assume normative behavior requires controlled processing, suggests that the dlPFC activates when prepotent responses conflict with desired normative outcomes^27, 28^. The neural activity of the anDDM, which arises from mutually inhibitory pools of option neurons receiving weighted inputs from hedonic and virtuous attributes, is in some ways consistent with such an interpretation. However, it calls into question assumptions that prepotency equates to hedonism, or even to automaticity^29^ more generally. Instead, our model suggests that the “prepotent response” may correspond, at least in the realm of value-based decision making, to choices consistent with the choice attribute that is currently receiving higher weight, *regardless of the source of that weight*. In other words, even when higher weights on normative attributes derive primarily from a deliberative, regulatory focus, as in our final two studies^23^, this results in *reduced* activity in the dlPFC when making normative choices (and greater activity when choosing hedonistically). Mechanistically, these patterns result from the fact that higher weights on normative attributes reduce the computation required for competitive neural interactions to settle on the normative response. Thus, while virtuous choices associated with successful self-control may sometimes recruit the posterior dlPFC, manipulations that increase the weight on normative attributes, either by making it more salient in the exogenous environment or focusing endogenous attention towards it, should both promote normative behavior and make it easier to accomplish.

This observation may help to explain why some researchers have found evidence consistent with greater response in the dlPFC promoting normative choice^1, 3, 6, 9, 30, 31^, while others have not^10–12^. Variations that influence the weight on normative attributes—whether across individuals, goal contexts, or paradigms—will tend to reduce statistical significance and increase heterogeneity in the link between neural activation in the dlPFC and normative choice. Fortunately, our model provides a way to predict both *when* and *why* dlPFC activity will be observed. For example, in the domain of intertemporal choice, our model predicts that making future outcomes more salient should amplify their weight in the choice process, promoting patience while *decreasing* dlPFC activation. This is exactly what is observed empirically^12^. Thus, researchers would do well to interpret activation of the dlPFC for a particular kind of choice (be it generous or selfish, healthy or unhealthy, patient or impatient) with caution. Such a pattern may say less about whether the dlPFC (and by extension, cognitive control more generally) is *required* to inhibit instinctual responses and preferences, and more about what kinds of attributes are most salient or valuable in the moment.

Our results have important implications for theories of self-control suggesting that the dlPFC promotes self-control by modulating attribute weights in the choice process^1, 31, 32^. The region of dlPFC that we observe here correlating with the anDDM is nearly identical to areas observed when dieters made healthy compared to unhealthy choices^1^, and when participants are required to recompute values based on contextual information^32^. Yet we find that the relationship between self-control “success” and “failure” in this region reverses when participants actively focus on health: dlPFC now responds more strongly to *unhealthy* choices. These results thus seem incongruent with the notion that this area down-regulates weight on norm-inconsistent considerations and up-regulates norm-consistent ones, since we observed *decreased* responses in this area in the context of *increased* normative choice and *increased* weight on normative attributes (Figure 4, c.f. ^23^). Moreover, we found no evidence that regulatory instructions led to greater overall activation in the dlPFC, as might be expected if this area implements changes in the weight given to normative attributes. Instead, this region appeared to correlate with the evidence accumulation stage of decisions, rather than with the evidence construction stage, responding during decision conflict generally, regardless of whether that conflict derived from greater weighting of hedonic or normative attributes.

We emphasize, however, that our results and conclusions apply narrowly to the area of dlPFC identified. The anDDM-related dlPFC region in this study lies posterior and dorsal to another dlPFC area that we have observed, in these same datasets, to track hedonistic and normative attributes in a goal-consistent manner and to serve as a candidate for mediating regulation-induced changes^23^. Furthermore, gray matter volume in this more anterior dlPFC area, but not in the posterior dlPFC region identified here, correlates with regulatory success^33^. Thus, while some areas of the dlPFC may indeed play an important role in promoting self-regulation and normative behavior by altering attribute weights in decision value, we suspect that they are anatomically and computationally distinct from the region of the posterior dlPFC sometimes assumed to serve this role. Future work will be needed to better delineate subregions of the dlPFC, and to determine the unique role each one plays in promoting normative choices.

The close correspondence between predictions of the anDDM and activation patterns in the dlPFC makes it tempting to conclude that this region performs this computational function. While this hypothesis is consistent with results from single-cell recordings^17, 34^, we also acknowledge that the dlPFC has been associated with many computational functions and roles, not all of which are mutually incompatible. Thus, it is possible that the dlPFC region observed here performs some sort of process that is correlated with, but not identical to, the neuronal computations of the anDDM.

Future work, including computational modifications or additions to the anDDM, as well as recordings from other modalities^34^, may help not only to elucidate the precise computational functions served by this area, but also the ways in which it promotes adaptive choice and normative behavior. Work extending these findings to other domains of normative choice, such as moral decision making or intertemporal choice, may also help to identify the commonalities and differences across different self-control dilemmas.

## Methods

### Computational Model Simulations

Our attribute-based neural drift diffusion model (anDDM: Figure 1) assumes that brain areas involved in decision making (particularly those that convert preferences into action) contain two spatially intermingled populations of neurons representing the options under consideration (here denoted as Option 1 and Option 2), with instantaneous firing rates (*FR*) at time *t* of *FR1(t)* and *FR2(t)*. At the beginning of the choice period *FR1*(0) = *FR2*(0) = 0. Firing rates in each population evolve dynamically from the onset of choice based on the sum total of excitatory and inhibitory inputs (detailed below). A choice results at time *t’*, the first moment at which the firing rate of one of the two populations exceeds a predetermined threshold or barrier *B*. The total response time RT is *t’* plus a constant non-decision time (*ndt*) that accounts for perceptual and motor delays.

Firing rates in the two pools evolve noisily over time according to the following two equations:

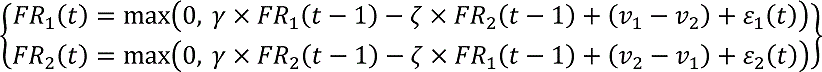

where the noise terms *ε_x_(t)* are normally distributed ∼N(0,.1), γ ≥ 1 represents recurrent auto-stimulation from the pool onto itself, ζ ≥ 0 represents inhibitory input from the other pool, and *v_1_* and *v_2_* represent external inputs proportional to the overall values of Options 1 and 2, determined by the weighted sum of their choice-relevant attribute values:

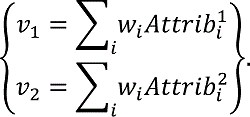

Thus, each pool’s activity receives an external input proportional to its value relative to the other option. In our simulations, we assumed two independent attributes: one related to hedonism (e.g., tastiness of a food) and one related to norms and standards (e.g. healthiness), although in principle any number and type of attribute could occur. Using these equations allowed us to simulate the dynamically evolving balance of excitation and inhibition across the two neuronal populations, and to derive distributions of both response times (RTs) and neural response. We label the final output (i.e., choice) of the system as “normative” if it results in selecting the option with the higher unweighted value for the normative attribute (e.g., the option with higher healthiness).

To simulate everyday self-control dilemmas using this framework, we simulated choices between two options representing different combinations of hedonistic and normative attributes, allowing the relative value difference between an option and its alternative on a given attribute to vary independently in the arbitrarily chosen range [−3, −2, … +2, +3]. This permitted us to explore how the likelihood of a normative choice changes depending on how much better or worse one of the two options is along hedonic and normative attribute dimensions, as well as what happens when the relative values of the two attributes conflict (i.e. take opposite signs) or do not.

We also sought to capture in our simulations the notion that a decision maker can vary from moment to moment in their commitment to and desire for hedonistic vs. normative goals. For example, a dieter may begin to relax the importance they place on norm-consistent attributes like healthiness once they reach their target weight, resulting in more unhealthy choices. In the main text (and Figure 2), we focus on simulations for two different goal contexts: one with a higher weight on tastiness, a hedonic attribute (i.e., *w_T_* = .05, *w_H_* = .02) and one with a higher weight on healthiness, a normative attribute (*w_T_* = .02, *w_H_* = .05). For simplicity, we assumed that all choices used a choice-determining threshold *B*=0.15, selected to produce RTs in the range typically observed in human subjects. Thus, for purposes of illustration, we simulated a decision-maker in two different contexts with different weights on the two attributes, facing 49 distinct choices representing different combinations of attribute values. To ensure that our conclusions held across a variety of weights, we also simulated an additional 34 different goal contexts, fully covering the factorial combination of weights on w_T_ and wH in the range of 0, .01, .02 … .05. Using these values and weights, we simulated choice frequencies, total neural activation (summed across the two neuronal pools), and RTs for each of the different hypothetical option pairs/attribute combinations, probing the effects of attribute weights, attribute magnitudes, and attribute conflict (i.e. match or mismatch between the signs of normative and hedonic attribute). Results of these simulations are displayed in Figure 2. Code is available at [link released after publication].

#### Experimental Studies

Details about portions of Studies 1, 2 and 3, as well as neuroimaging parameters, have been reported previously^22, 23^. Here, we highlight in brief the most important details for the current work.

##### Participants

For Study 1, we analyzed data from 51 male volunteers (mean age 22, range 18-35). All participants received a show-up fee of $30 as well as an additional amount ranging from $0-$100, depending on the outcome of the task (see below). For Study 2, we analyzed data from 49 volunteers (26 male, mean age 28, range 19-40). For Study 3, 36 individuals from Study 2 returned to the lab for a separate session on a separate day to complete a dietary choice task. For each session in Studies 2 and 3, participants received a show-up fee of $50. Participants completing the altruistic choice task in Study 2 also received from $0-$40 in additional earnings, depending on the outcome of the task (see below). Caltech’s Internal Review Board approved all procedures. Participants in all studies provided informed consent prior to participation.

#### Tasks and Stimuli

##### Altruistic Choice Task (Studies 1 & 2)

We examined self-control dilemmas pitting self-interest against generosity using an Altruistic Choice Task for Studies 1 and 2. On every trial in the scanner, the participant chose between a proposed pair of monetary prizes to herself and a real but anonymous partner, or a constant default prize-pair to both ($50 in Study 1, $20 in Study 2) (Figure 3a-b). Proposed prizes in the prize-pair varied from $0 to $100 in Study 1 and $0 to $40 in Study 2, and always involved one individual receiving an amount less than or equal to the default, while the other individual received more. Thus, on every trial the participant had to choose between generous behavior (benefitting the other at a cost to oneself) and selfish behavior (benefitting oneself at a cost to the other).

Upon presentation of the proposal, participants had up to four seconds to indicate their choice using a 4-point scale (Strong No, No, Yes, Strong Yes), allowing us to simultaneously measure both their decision and strength of preference at the time of choice. The direction of increasing preference (right-to-left or left-to-right) varied for each round of the task in Study 1, and across participants in Study 2. If the subject did not respond within four seconds, both individuals received $0 for that trial.

To increase the anonymity of choices, the participant’s choice was implemented probabilistically: in 60% of trials he received his chosen option, while in 40% of trials his choice was reversed and he received the alternative, non-chosen option. This reversal meant that while it was always in the participant’s best interest to choose according to her true preferences, her partner could never be sure about the actual choice made. Probabilistic implementation does not strongly influence the choices participants make^22, 23^, but permits more plausible anonymity, increasing the self-control challenge involved in choosing generously. The participants were informed that the passive partners were aware of the probabilistic implementation, and the outcome was revealed on every trial 2-4 seconds following the response.

Study 1 included 180 trials total, with no specific instructions for how to respond. Study 2 included 270 trials, 90 each in three instructed focus conditions. See the *Manipulating Normative Goals (Studies 2 & 3)* section below for details on these instructions.

##### Dietary Choice Task (Study 3)

We examined self-control dilemmas in a second context pitting hedonism against healthy eating using a Dietary Choice Task for Study 3. Prior to the task, participants rated a set of 200 different foods for their healthiness and tastiness. These ratings were used to 1) select a pool of 90 foods that covered a range of health and taste ratings and 2) select a neutral reference food rated as neutral on both health and taste.

On each of 270 trials in the scanner, participants saw one of the 90 different pre-selected foods (Figure 3c), and had to decide whether they would prefer to eat the displayed food or the reference food. As in the altruistic choice task, participants had up to four seconds to indicate their choice using a 4-point scale (Strong No, No, Yes, Strong Yes). If the subject did not respond within four seconds, one of the foods was selected randomly. To match the instructed attention manipulation used in the Altruistic Choice Task, participants completed 90 trials each in one of three instructed focus conditions. See the *Manipulating Normative Goals (Studies 2 & 3)* section below for details.

To match the probabilistic outcome used in the altruistic choice task, the participant’s choice was also implemented probabilistically in the Food Choice Task. In 60% of trials he received his chosen option, while in 40% of trials his choice was reversed and he received the alternative, non-chosen option. To reduce the length of the task, participants did not see this outcome on every trial. Instead, three trials were selected randomly at the end of each scan, and participants viewed their choice as well as the probabilistic outcome on that trial.

##### Manipulating Normative Goals (Studies 2 & 3)

Our computational model simulations suggested that the extent to which normative choices are associated with greater neural response depends to a large extent on the priority or weight given to normative vs. hedonic attributes. We thus capitalized on the design of Studies 2 and 3, which manipulated attention to different attributes (and corresponding weights), allowing us to test specific predictions of the anDDM.

##### Generosity Manipulation (Study 2)

To manipulate attention to different attributes, during the Altruistic Choice Task in Study 2, participants completed trials in one of three different instructed focus conditions: Respond Naturally, Focus on Ethics, and Focus on Partner. During *Natural* trials, participants were told to allow whatever feelings and thoughts came most naturally to mind, and to just choose according to their preferences on that trial. During *Ethics* trials, participants were asked to focus on doing the right thing during their choices. They were encouraged to think about the justice of their choice, as well as its ethical or moral implications, and to try to bring their actions in line with these considerations. During *Partner* trials, participants were asked to focus on their partner’s feelings during their choices. They were encouraged to think about how the other person would be affected, as well as whether they would be happy with the choice, and to bring their actions in line with these considerations.

Each participant completed 90 trials per condition, presented in randomly interleaved blocks of ten trials. A detailed set of instructions informing participants of their task for the upcoming block of trials was presented for 4 seconds prior to the block, and participants were asked to focus on the specific instruction for all trials within that block.

##### Healthiness Manipulation (Study 3)

Analogous to the Altruistic Choice Task in Study 2, we manipulated healthy eating in Study 3 using an instructed focus manipulation. Each participant completed 270 choice trials, 90 each in one of three attentional conditions: *Natural* Focus, *Taste* Focus, or *Health* Focus. During *Natural* trials, participants were told to allow whatever feelings and thoughts came most naturally to mind, and to just choose according to their preferences on that trial. During *Taste* trials, participants were asked to focus on how tasty each food was, and to try to bring their actions in line with this consideration. During *Health* trials, participants were asked to focus on the health implications of their choice. As in the Altruistic Choice Task, attentional instructions were given prior to each block of 10 trials, and participants were asked to focus on the specific instruction given for all trials within a block. However, participants knew that they would receive the outcome of one of their choices, and were told that they should choose according to their preferences regardless of the instruction, thus encouraging participants to choose in a way that reflected their current decision value for the item.

#### Defining Normative Choice

##### Behavioral definition of generosity

All choices involved a tradeoff between maximizing outcomes for the self or for the other. We therefore label specific decisions as normative (i.e., generous) if the participant accepted a proposal when $Self < $Other, or rejected one when $Self > $Other. Choices were labeled as hedonistic (i.e., selfish) otherwise.

##### Behavioral definition of healthy choice

In the Dietary Choice Task, we separately examined trials requiring a tradeoff between taste and health (i.e. conflict trials where a food was rated either as healthy but not tasty, or as unhealthy but tasty) as well as trials with no tradeoff (i.e., no-conflict trials where a food was both tasty and healthy, or both unhealthy and not tasty). In both cases, we label specific decisions as normative (i.e., healthy) if the participant either accepted a healthy food, or rejected an unhealthy food. All other choices were labeled as hedonistic (i.e., unhealthy).

##### Computational Model Fitting

We used a Bayesian model-fitting approach to identify best-fitting model parameters of the anDDM (i.e. attribute weight parameters, threshold *B*, non-decision time *ndt*, auto-excitation parameter γ and lateral inhibition parameter ζ) to account for choices and RTs, separately for each participant in each study and (in Studies 2 and 3) each condition. More specifically, we obtained estimates of the posterior distribution of each parameter using the Differentially-Evolving Monte-Carlo Markov Chain (DEMCMC) sampling method and MATLAB^35^ code developed by ^36^. This method uses the anDDM described above (Computational Model Simulations) to simulate the likelihood of the observed data (i.e. choices and RTs) given a specific combination of parameters, and then uses this likelihood to construct a Bayesian estimate of the posterior distribution of the likelihood of the parameters given the data.

For each individual fit, we used 3 x *N* chains, where *N* is the number of free parameters (7 in Studies 1 and 2, 6 in Study 3), using uninformative priors and constraining parameter values as shown in Supplementary Table S1 based on previous work^22, 23^ and theoretical bounds. To construct the estimated posterior distributions of each parameter, we sampled 1500 iterations per chain after an initial burn-in period of 500 samples. Best-fitting values of each parameter were computed as the mean over the posterior distribution for that parameter. These parameter values (see Supplementary Table S1) were used to simulate trial-by-trial activation across the two neuronal pools for use in the GLMs described below. Importantly, parameter values identified by this fitting procedure suggested that the model provided a good fit to behavior across all three studies (Supplementary Figure 2).

#### Neuroimaging Analyses

##### GLM 1a: Correlates of the anDDM (Study 1)

We used GLM 1a to identify brain regions where activation varied parametrically according to the predictions of the anDDM in Study 1 (Altruistic Choice Task). To this end, we determined that the best BOLD approximation of the anDDM was a parametric modulator with a value consisting of the sum total of the simulated response across both pools of neurons, averaged over all simulations terminating in the observed choice on that trial within ±250ms of the observed RT, and modulating a boxcar function with onset at the beginning of the choice period and having a duration of the RT on that trial (see Supplemental Methods for further detail on selecting the best regressor). To simulate expected anDDM activation on each trial, we generated 5000 simulations using the best-fitting parameters for each participant and the estimated value of the proposal and default on each trial (i.e., w_Self_*$Self + w_Other_*$Other + w_Fairness_*|$Self - $Other|).

Then, for each subject we estimated a GLM with AR(1) and the following regressors of interest: R1) A boxcar function for the choice period on all trials (duration = RT on that trial). R2) R1 modulated by the subject’s stated preference on that trial (1 = Strong No, 4 = Strong Yes). R3) R1 modulated by the estimated activation of the anDDM on that trial. R4) A boxcar function of 3 seconds specifying the outcome period on each trial. R5) R4 modulated by the outcome for the self on each trial. R6) R4 modulated by the outcome for the partner on each trial. R7) A boxcar function (duration = 4 seconds) specifying missed trials. Parametric modulators were orthogonalized to each other in SPM. Regressors of non-interest included six motion regressors as well as session constants.

We then computed subject-level contrasts of the anDDM parametric modulator (R3) against an implicit baseline. Finally, to test the hypothesis that anDDM responses might correlate with activation in the dlPFC, we subjected this contrast to a one-sample t-test against zero, thresholded at a voxel-wise *P* < .001, and a cluster-defining threshold of *P* < .05, small-volume corrected within a 10-mm spherical region of interest (ROI) centered on the peak coordinates of activity for the contrast of normative (healthy) vs. hedonistic (unhealthy) choice in a previous study of self-control in dieters^1^. In addition to this ROI-analysis, we performed supplemental analyses at the whole-brain level at a voxel-level threshold of *P* < .001 uncorrected and a whole-brain cluster-corrected level of P < .05.

##### GLM 1b: Correlates of the anDDM (Study 2)

GLM1b was similar to GLM1a, with the exception that we estimated regressors for each condition separately. R1, R4, and R7 were boxcar functions representing the choice period for the *Natural*, *Ethics*, and *Partner* conditions, respectively. R2, R5, and R9 modulated R1, R4 and R7 with the decision value on that trial. R3, R6, and R9 modulated R1, R4, and R7 using the estimated activation of the anDDM on that trial. A single contrast representing neural correlates of the anDDM was constructed by combining R3, R6 and R9 at the subject-level and performing a one-sample t-test against zero, thresholded at a voxel-wise P < .001 and a small-volume cluster-corrected level of *P* < .05 within the dlPFC ROI described above.

##### GLM1c: Correlates of the anDDM (Study 3)

GLM1c was similar to GLM1b, but applied to the Food Choice Task. R1, R4, and R7 were boxcar functions representing *Natural*, *Taste*, and *Health* focus conditions. R2, R5, and R8 were parametric modulators representing the decision value on that trial, and R3, R6, and R9 were modulators consisting of anDDM activity simulated using healthiness and tastiness ratings as attributes. Similar to Studies 1 and 2, correlates of the anDDM were identified in this study thresholded at a voxel-wise P < .001 and a small-volume cluster-corrected level of *P* < .05 within the dlPFC ROI described above.

##### Data-driven ROI definition

Based on GLMs 1a, b and c, we identified a region of the left dlPFC consistently associated with the anDDM across all three studies through a three-way conjunction analysis using the imcalc function in SPM12, with each individual study map thresholded at P < .05, small-volume corrected, and a minimum overlap of > 5 contiguous voxels. Outside of this ROI, we also identified regions significant across all three studies at P < .05, whole-brain corrected. This identified just three regions, located in the left dlPFC, left IFG, and dACC (Figure 4 and Supplemental Figures S3 and S4). We then interrogated activation within these regions specifically for the contrast of normative vs. hedonistic choice, using GLMs 2a, b and c, as specified below.

##### GLM 2a: Generous vs. Selfish decisions in Altruistic Choice (Study 1)

We used GLM 2a to test predictions about activation on trials in which subjects chose generously or selfishly. The analysis was carried out in three steps.

First, for each subject we estimated a GLM with AR(1) and the following regressors of interest: R1) A boxcar function for the choice period on trials when the subject chose selfishly. R2) R1 modulated by the value of 4-point preference response (i.e., Strong No to Strong Yes) at the time of choice. R3) A boxcar function for the choice period on trials when the subject chose generously. R4) R3 modulated by behavioral preference. Regressors of non-interest included six motion regressors as well as session constants.

Second, we computed the subject-level contrast image [R3 – R1], which identified regions with differential response for generous compared to selfish choices. Seven subjects were excluded from this analysis for having fewer than 4 generous choices over the 180 trials. We computed the average value of this contrast within the three anDDM ROIs specified above. As a supplementary analysis, we also asked whether any voxels beyond these regions demonstrated a significant effect, using a whole-brain analysis thresholded at P < .001, uncorrected (see Supplementary Table S3).

##### GLM 2b: Generous vs. Selfish decisions in Altruistic Choice (Study 2)

We used GLM 2b to test predictions about activation on trials in which the subject chose generously or selfishly in Study 2, and to compare how instructed attention altered these responses. All unreported details are as in GLM1a. Regressors of interest consisted of the following: R1) A boxcar function for the choice period on trials when the subject chose selfishly in *Natural Focus* trials. R2) R1 modulated by the value of 4-point preference response (i.e., Strong No to Strong Yes) expressed at the time of choice. R3) A boxcar function for the choice period on trials when the subject chose generously in *Natural Focus* trials. R4) R3 modulated by behavioral preference. R5-R8) Analogous regressors for generous and selfish choices during *Ethics Focus* trials. R9-12) Analogous regressors for generous and selfish choices during *Partner Focus* trials. R13-15) A boxcar function of 3 sec duration signaling the outcome period for *Natural, Ethics,* or *Partner Focus* trials. R16-18) R13-15 modulated by the amount received by the subject at outcome. R19-21) R13-15 modulated by the amount received by the partner at outcome.

We then computed the subject-level contrast images [R3 – R1], [R7 – R5], and [R11 – R9], which identified regions with differential response for generous compared to selfish choices in each condition. We computed the average value of each of these contrasts within the three anDDM ROIs specified above. As a supplementary analysis, we also asked whether any voxels beyond these regions demonstrated a significant effect in any condition, using a whole-brain analysis thresholded at P < .001, uncorrected (see Supplementary Table S3).

##### GLM 2c: Healthy vs. Unhealthy decisions in the Food Choice Task (Study 3)

GLM 2c was analogous to GLM 2b, but examined healthy vs. unhealthy choices in the Dietary Choice Task, separately for conflicted trials (i.e. healthy but not tasty foods and tasty but unhealthy foods) and for unconflicted trials (i.e. healthy and tasty foods or unhealthy and not tasty foods). It included the following regressors of interest: R1) A boxcar function for the choice period on conflicted trials when the subject made a healthy choice (i.e., accepted a healthy-but-not-tasty or rejected a tasty-but-unhealthy food) in *Natural Focus* trials. R2) R1 modulated by the value of behaviorally expressed preference at the time of choice. R3) A boxcar function for the choice period on conflicted trials when the subject made an unhealthy choice in *Natural* trials. R4) R3 modulated by behavioral preference. R5-8) Analogous regressors for healthy and unhealthy choices during conflicted *Taste Focus* trials. R9-12) Analogous regressors for healthy and unhealthy choices during *Health Focus* trials. R13) Healthy choices on unconflicted *Natural Focus* trials. R14) Unhealthy choices on unconflicted *Natural Focus* trials. R15-16) R13 and R14 modulated by preference. R17-R20) Analogous regressors for healthy and unhealthy choice on unconflicted trials in the *Health Focus* trials. R21-R24) Analogous regressors for healthy and unhealthy choice on unconflicted trials in the *Taste Focus* trials. Subject-level contrast images of healthy vs. unhealthy choices, in each condition separately and separately for conflicted vs. unconflicted trials, were computed in a manner identical to GLM2b. We analyzed activation for these contrasts specifically within the three ROIs identified as anDDM regions. As a supplementary analysis, we also report results at the whole-brain level at P < .001, uncorrected, in Table S3. Unreported details are as in GLM 2a.

## Data Availability

Behavioral data and all analysis code are available on the Open Science Framework at [link released after acceptance for publication]. Neuroimaging data are available upon request to the authors.

## Acknowledgments

These studies were made possibly by grants from the Gordon and Betty Moore Foundation, the Lipper Foundation, and a National Institute of Mental Health Silvio O. Conte award (NIMH Conte Center 2P50 MH094258). The scientific results and conclusions reflect the authors’ opinions and not the views of the granting entities. We gratefully acknowledge Antonio Rangel for comments on earlier versions of this manuscript.

## Author Contributions

All authors contributed to the design of the studies. C.H. and A.T. collected the data, and C.H. analyzed the data and developed the computational model. C.H., and A.T. wrote the paper.

## Supplementary Materials

### Supplementary Methods

#### Choosing the appropriate fMRI regressor for the anDDM model (GLMs 1a, b and c)

The attribute-based neural drift diffusion model (anDDM) produces a dynamic accumulation signal that builds over hundreds of milliseconds. This raises a question about the appropriate way to model this signal in the hemodynamic response, which evolves more slowly over 5-10 seconds. To determine the appropriate regressor for GLMs 1a, b, and c, we simulated 5000 instantiations of the anDDM for every subject and trial in Study 2, using a time step of 5 ms. For each subject, we then averaged the 5000 simulations at each time point to produce a single time course of total activity across the two neuronal pools for a given set of trials. We convolved this simulated time course with the canonical form of the hemodynamic response function (HRF) to construct an expected BOLD time series given the inputs. We refer to this as the *ideal BOLD*. We then compared the shape of the ideal BOLD to two different possible instantiations within a traditional GLM analysis in SPM. Version 1 consisted of a parametric modulator of a stick function placed at the onset of the trial, consisting of the sum total activity in the anDDM for each trial, *∑^RT^_t=1_ FR_1_(t) + FR_2_(t)*. Version 2 consisted of a parametric modulator identical to Version 1, but modulating a boxcar function placed at the onset of the trial with duration equal to RT for that trial. Each of these regressors was convolved with the canonical form of the HRF and correlated with the ideal time series to determine the one providing the closest match.

Results suggested that version 2 provided a closer match (Pearson’s *r* ranging from .90-.99, average = .96) compared to version 1 (Pearson’s *r* ranging from .62-.94, average = .82). Note also that the inclusion of the unmodulated boxcar function with duration equal to the RT on each trial controls for non-specific activation related to response times that does not build over time in the manner expected based on the anDDM.

### Supplementary Results

In the main paper, we focus on the effects of normative vs. hedonistic choice within the dlPFC ROI defined by the conjunction of anDDM-correlated trial-by-trial activity across all three studies. However, in addition to this dlPFC ROI, we identified two other regions, in the dorsal anterior cingulate cortex (dACC, see Figure S3) and left inferior frontal gyrus (IFG)/anterior insula (IFG/aIns, see Figure S4) whose activity correlated with the anDDM across all three studies (P < .001, whole brain corrected within each study). Here, we report analogous results on measures of BOLD response in these regions during normative vs. hedonistic choice, for the sake of completeness. These results suggest that our results are a general principle of areas correlating with anDDM response.

### dACC response during normative vs. hedonistic choices in Studies 1, 2, and 3

We began by examining whether activity in the dACC correlated with the contrast of normative (generous) vs. hedonistic (selfish) choices in Study 1. As expected, and similar to the dlPFC, this region showed a significantly greater response during generous compared to selfish choices (paired t_43_ = 3.4825, *P* = .001, Figure S3d). Similarly, in Study 2, we observed a significant effect of normative goals on the difference in response between normative and hedonistic choices (*F*_2,96_ = 13.67, *P* = 5.97 × 10^-6^). Follow-up t-tests confirmed that this was driven by a stronger response in the dACC to normative (generous) choices in Natural trials (paired-t_43_ = 3.53, *P* = .0009) as well as significantly stronger response to *hedonistic* choices (paired-t_43_ = 2.41, *P* = .02) during Partner-focused trials. Finally, we replicated a similar pattern of effects in Study 3, showing a significant influence of normative (i.e., health-focused) goals on the contrast of normative vs. hedonistic choices (*F*_2,96_ = 3.64, *P* = .03), which was driven by a stronger response on normative (healthy) choices in the Natural and Taste conditions, and a marginally stronger response on *hedonistic* (i.e., unhealthy) choices during Health Focus trials (paired-t_43_ = 1.96, *P* = .058).

### IFG/aIns response during normative vs. hedonistic choices in Studies 1, 2, and 3

As expected if IFG/aIns response correlates with the anDDM, we observed similar patterns of responding on normative vs. hedonistic choices across all three studies within this region.

IFG/aIns showed a significantly greater response during generous compared to selfish choices (paired t_43_ = 3.22, *P* = .002, Figure S4d). Similarly, in Study 2, we observed a significant effect of normative goals on the difference in response between normative and hedonistic choices (*F*_2,96_ = 17..66, *P* = 2.93 × 10^-7^, Figure S4e). Follow-up t-tests confirmed that this was driven by a stronger response in the dACC to normative (generous) choices in Natural trials (paired-t_43_ = 5.06, *P* = 6.57× 10^-6^) as well as significantly stronger response to *hedonistic* (i.e., selfish) choices (paired-t32 = 2.66, *P* = .01) during Partner-focused trials. Finally, we replicated a similar though non-significant pattern of the effects of normative goals in Study 3 (*F*_2,96_ = .75, *P* = .39, Figure S4f). However, planned post-hoc comparisons confirmed that activation in the left IFG/aIns was stronger on normative (healthy) choices in the Natural condition (paired-t_43_ = 2.65, *P* = .01), while activation for this same condition was non-significantly reversed on Health Focus trials (*P* = .66). The direct comparison of normative vs. hedonistic choices during Natural vs. Health Focus was also significant (paired-t34 = 2.18, *P* = .04).

## Supplementary Figures

**Figure S1.**
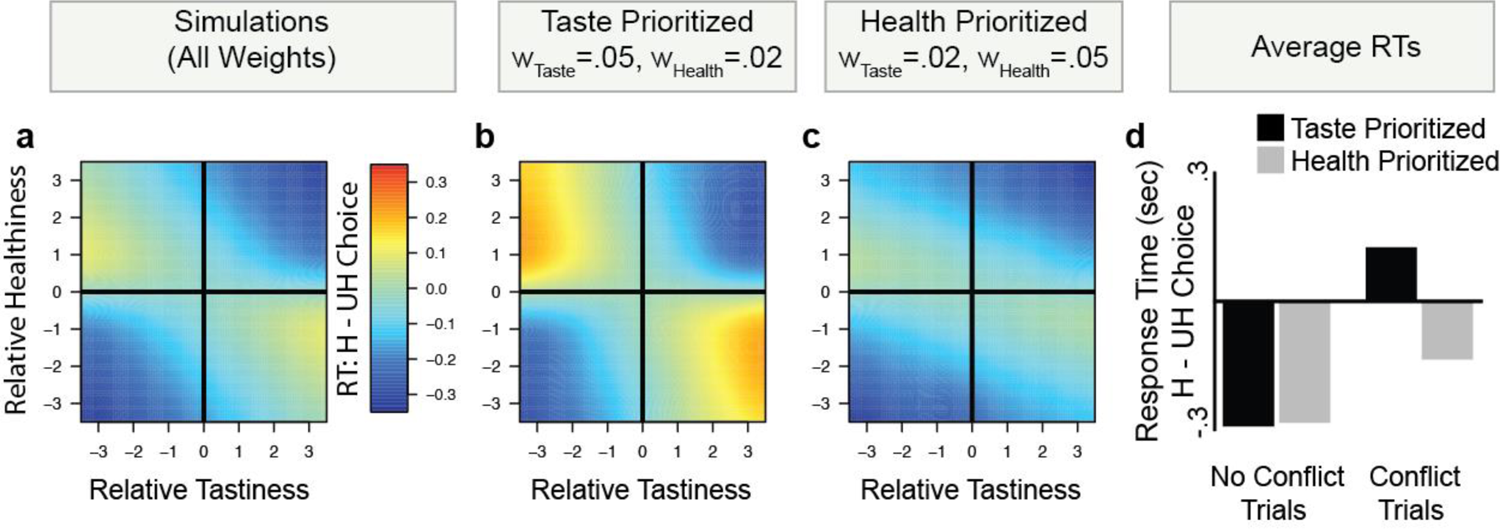
Computational simulations of response time (RT). **(a)** Similar to neural response, model simulations suggest that response times when making normative (i.e., healthy, H) choices instead of hedonistic (i.e. unhealthy, UH) ones (i.e., RTH – RTUH) depends on relative healthiness and tastiness for goal contexts that prioritize both **(b)** hedonism and **(c)** normative goals. Warmer colors indicate longer RTs for healthy choices, indicated by larger differences in RTH – RTUH. **(d)** Average differences in RT for health compared to unhealthy choices (averaging over different options with different attribute values) are displayed for contexts in which health or taste are prioritized, divided as a function of whether relative healthiness and tastiness conflict (i.e., take opposite signs) or do not (no conflict trials). In no conflict trials, on average, healthy choices are easy regardless of whether taste is prioritized (black bars) or health is prioritized (gray bars), indicated by comparatively faster RTH than RTUH. In conflict trials, however, on average, healthy choices are difficult only in when taste is prioritized (when w_Taste_ > w_Health_), reflected in relatively longer RTH than RTUH.

**Figure S2.**
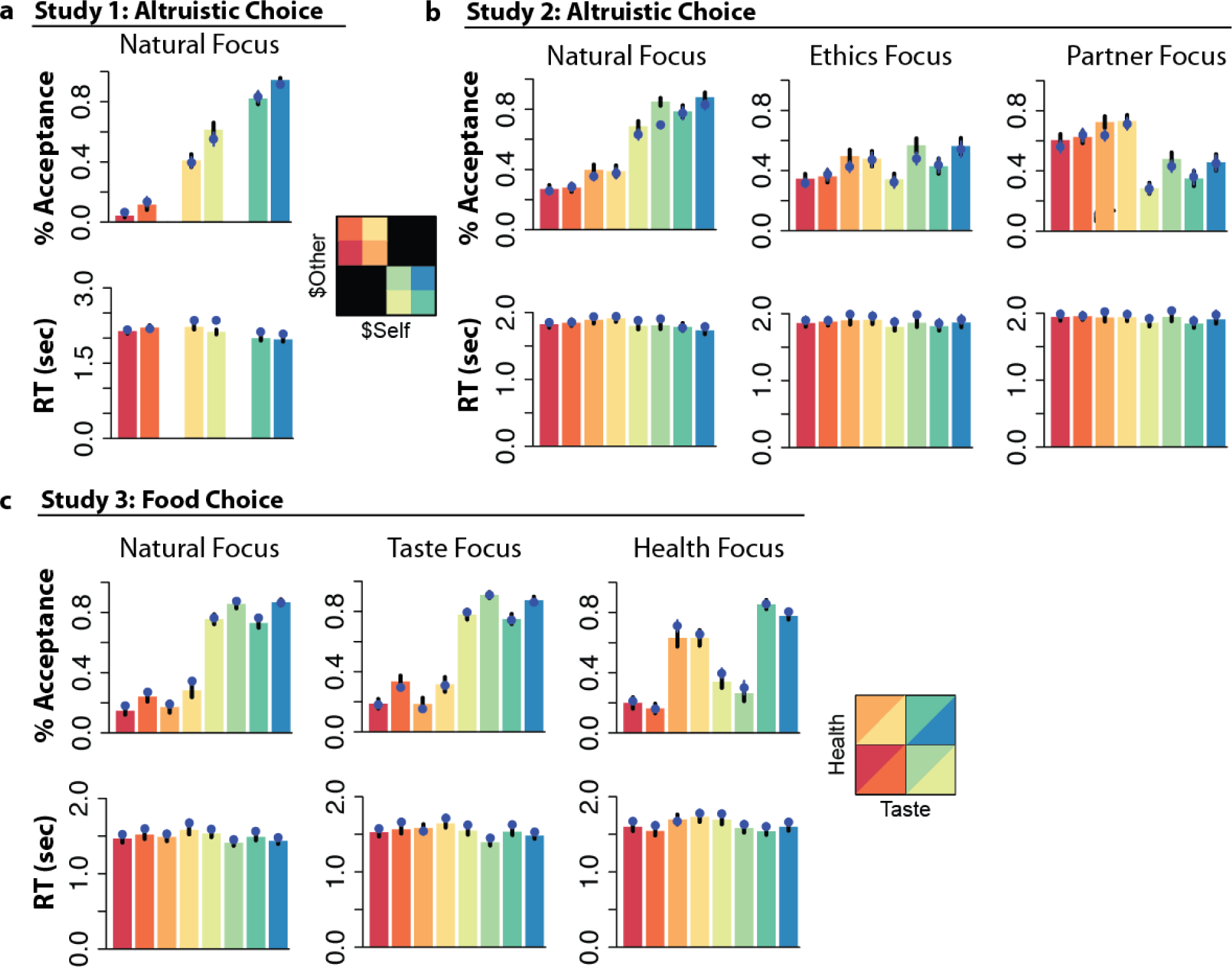
Model fits to behavior. (**a**) Choices and RTs for observed behavior (colored bars) and model simulations (blue dots) for different choice types in Study 1. (**b**) Observed and model-simulated choices and RTs in Study 2, separately by regulatory condition. (**c**) Observed and model-simulated choices and RTs in Study 3, separately by regulatory condition. Error bars show standard error of the mean.

**Figure S3.**
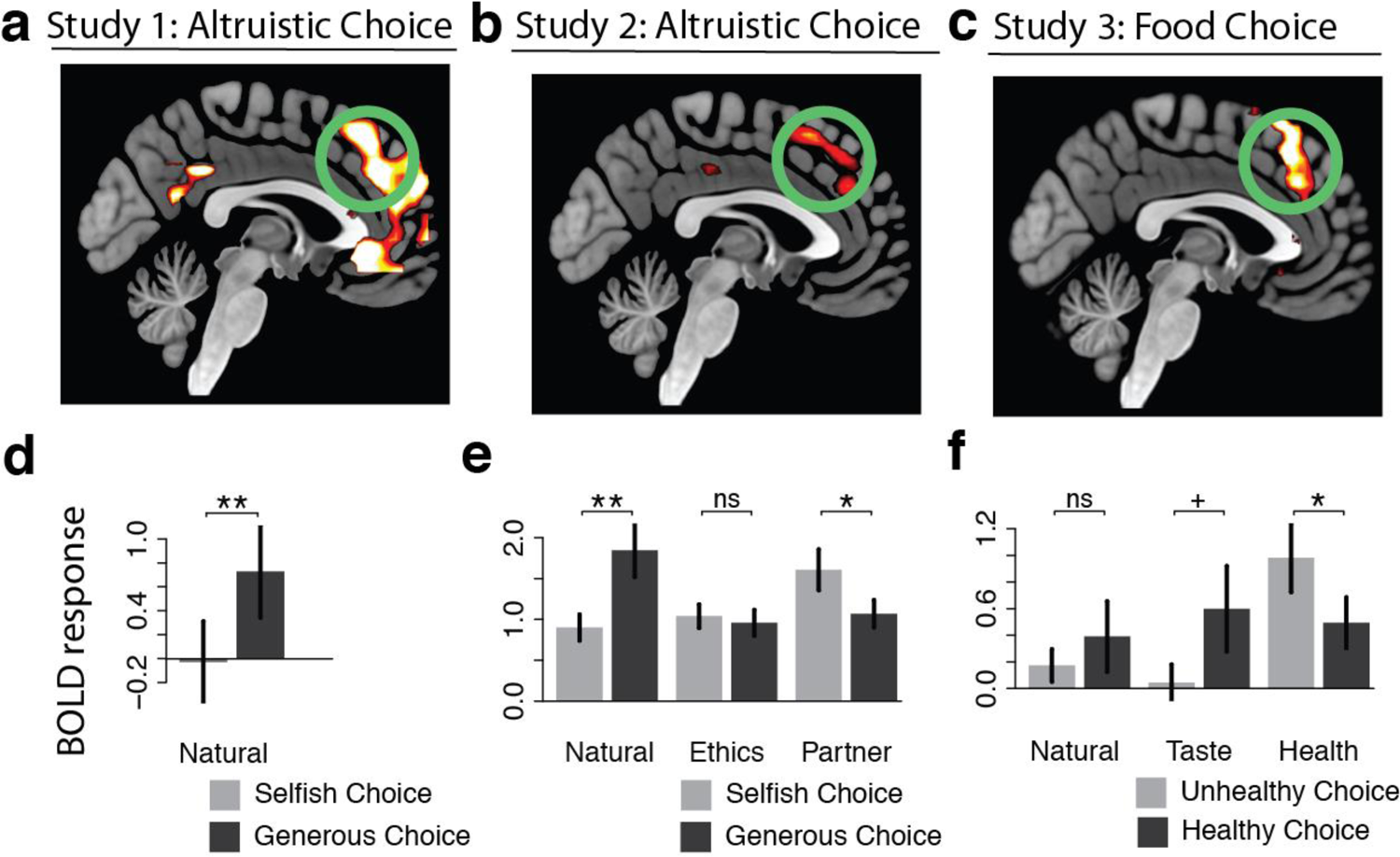
BOLD responses in the anterior cingulate cortex during self-control dilemmas. Top: Trial-by-trial BOLD response in the dACC correlates with predicted activity of the anDDM across three separate studies, including during both altruistic choice **(a, b)** and during dietary choice **(c)**. All maps thresholded at *P* < .001 uncorrected for display purposes. Bottom: Within the dACC ROI defined by the three-way conjunction of anDDM response across all studies, BOLD response during normative choice (black) vs. hedonistic choice (light gray) when attributes conflict, in **d**) Study 1 for all trials, as well as in **e**) Study 2 and **f**) Study 3 as a function of regulatory goals. As predicted, normative choices activate the dACC, but only when goals result in a greater weight on hedonistic than normative attributes. + *P* < .05, one-tailed; * *P* < .05; ** *P <* .01.

**Figure S4.**
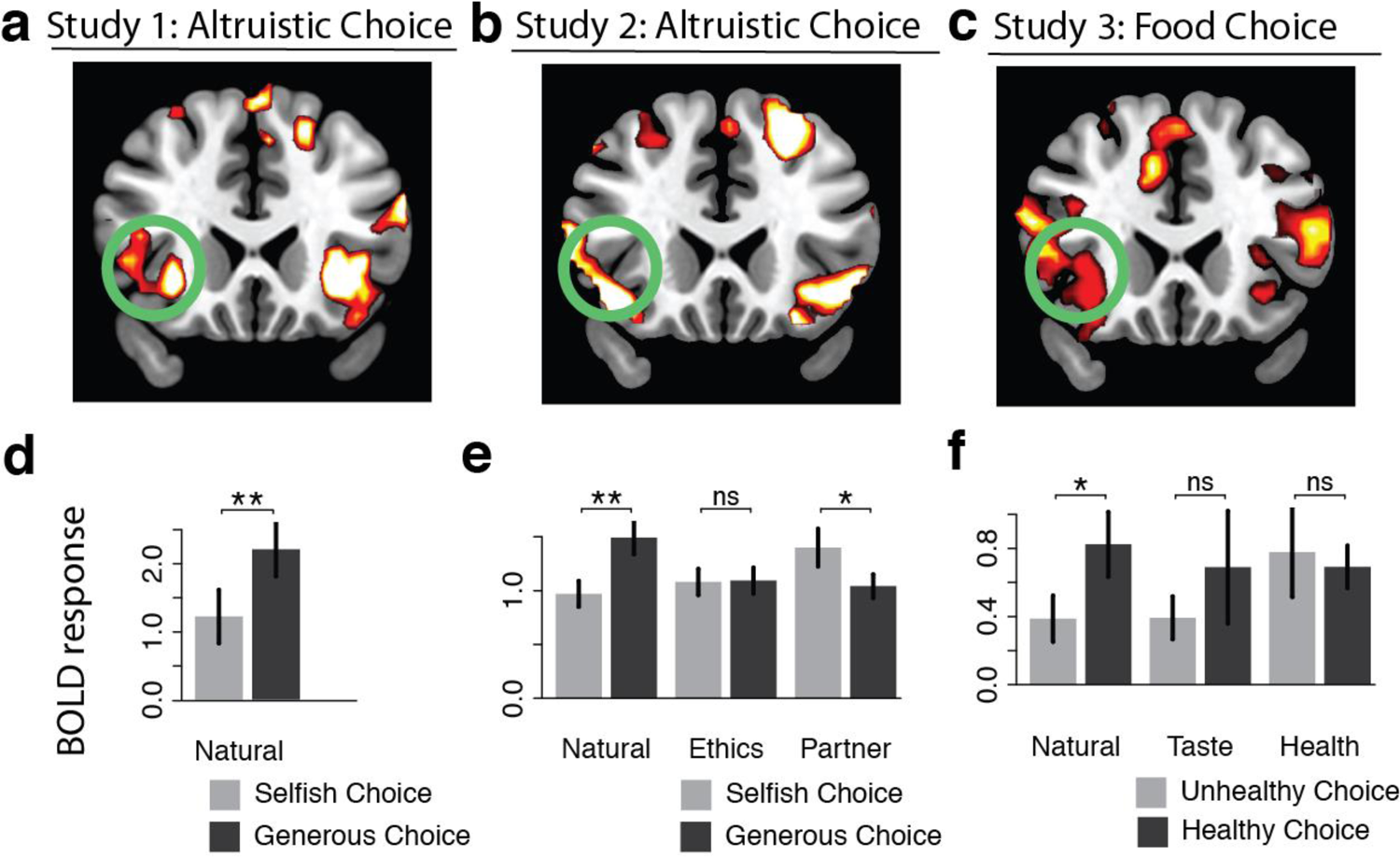
BOLD responses in the inferior frontal gyrus (IFG)/anterior insular cortex during self-control dilemmas. Top: Trial-by-trial BOLD response in the IFG/insula correlates with predicted activity of the anDDM across three separate studies, including during both altruistic choice **(a, b)** and during dietary choice **(c)**. All maps thresholded at *P* < .001 uncorrected for display purposes. Bottom: Within the IFG/insula ROI defined by the three-way conjunction of anDDM response across all studies, BOLD response during normative choice (black) vs. hedonistic choice (light gray) when attributes conflict, in **d**) Study 1 for all trials, as well as in **e**) Study 2 and **f**) Study 3 as a function of regulatory goals. As predicted, normative choices activate the IFG/insulas, but only when goals result in a greater weight on hedonistic than normative attributes. + *P* < .05, one-tailed; * *P* < .05; ** *P <* .01.

**Table S1.** Estimated Model Parameters

| Parameter | *A priori*<br>constraints | Study 1 | Study 2,<br>Natural | Study 2, Ethics | Study 2,<br>Partner | Study 3,<br>Natural | Study 3, Taste | Study 3,<br>Health |
| --- | --- | --- | --- | --- | --- | --- | --- | --- |
| *w<sub>Self</sub>* | -.5 to +.5 | .0036±.0011 | .0073±.0035<sup>a</sup> | .0061±.0047<sup>a</sup> | .0037±.0065<sup>b</sup> | - | - | - |
| *w<sub>Other</sub>* | -.5 to +.5 | .0008±.0015 | .001±.0038<sup>a</sup> | .0041±.0045<sup>b</sup> | .0051±.0038<sup>b</sup> | - | - | - |
| *w<sub>Fairness</sub>* | -.5 to +.5 | .0008±.001 | .0017±.0033<sup>a</sup> | .0053±.0046<sup>b</sup> | .0024±.0035<sup>a</sup> | - | - | - |
| *w<sub>Taste</sub>* | -.5 to +.5 | - | - | - | - | .0074±.0027<sup>a</sup> | .0077±.0029<sup>a</sup> | .002±.0028<sup>b</sup> |
| *w<sub>Health</sub>* | -.5 to +.5 | - | - | - | - | -.0002±.0018<sup>a</sup> | -.0008±.0018<sup>a</sup> | .0055±.0034<sup>b</sup> |
| *B* | 0 to +1.0 | .3181±.1425 | .2773±.1373<sup>a</sup> | .3628±.1453<sup>b</sup> | .4062±.1586<sup>b</sup> | .1691±.0501<sup>a</sup> | .1821±.0616<sup>a,b</sup> | .2009±.0819<sup>b</sup> |
| *ndt* | 0 to +2.0s | .8002±.215 | .5989±.2219<sup>a</sup> | .4835±.1448<sup>b</sup> | .4859±.1322<sup>b</sup> | .5442±.1321<sup>a</sup> | .5397±.1361<sup>a</sup> | .5399±.1589<sup>a</sup> |
| $\zeta$ | 0 to +2.0 | .583±.3034 | .5531±.3086<sup>a</sup> | .7469±.2814<sup>b</sup> | .7592±.2806<sup>b</sup> | .4102±.0768<sup>a</sup> | .4093±.0865<sup>a</sup> | .3958±.0848<sup>a</sup> |
| $\gamma$ | +1.0 to +3.0 | 1.8979±.3575 | 2.0148±.3881<sup>a</sup> | 2.2043±.3744<sup>a,b</sup> | 2.2952±.3467<sup>b</sup> | 1.6435±.1279<sup>a</sup> | 1.654±.1578<sup>a</sup> | 1.6802±.1455<sup>a</sup> |

*Note.* Parameter values were estimated using a Differential-Evolution Markov Chain Monte Carlo method developed by Holmes and Trueblood<sup>1</sup>. Parameters beginning with *w* indicate weighting parameters applied to different attributes (Studies 1 and 2: proposed payoff to self vs. the default, proposed payoff to other vs. the default, and fairness [ $\$Self - \$Other$ ]; Study 3: tastiness and healthiness vs. the default). *B*: choice-defining threshold. *ndt*: non-decision time. $\zeta$ : lateral inhibition parameter from one neuronal pool onto the other. $\gamma$ : auto-excitation parameter from a neuronal pool onto itself. *A priori* constraints on the parameters, determined based on previous work and on theoretical limits, restricted them to the range indicated. In Studies 2 and 3, columns indicated by different subscripts differ significantly from each other at $P < .05$ , corrected for multiple comparisons.

**Table S2.**
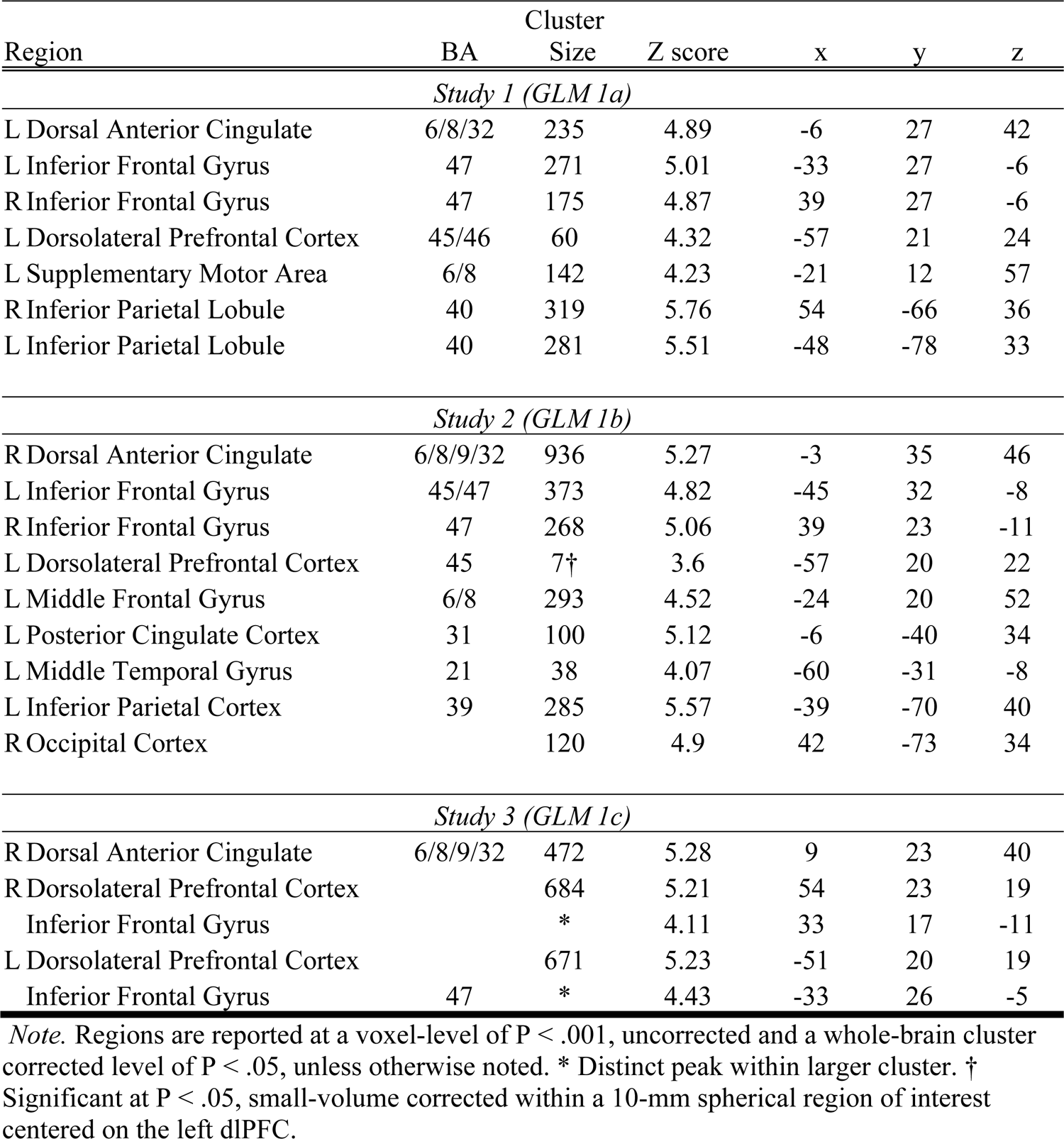
Neural correlates of the attribute-based neural drift diffusion model across <u>studies</u>

| Region | Cluster |  | Z score | x | y | z |
| --- | --- | --- | --- | --- | --- | --- |
|  | BA | Size |  |  |  |  |
| Study 1 (GLM 1a) |  |  |  |  |  |  |
| L Dorsal Anterior Cingulate | 6/8/32 | 235 | 4.89 | -6 | 27 | 42 |
| L Inferior Frontal Gyrus | 47 | 271 | 5.01 | -33 | 27 | -6 |
| R Inferior Frontal Gyrus | 47 | 175 | 4.87 | 39 | 27 | -6 |
| L Dorsolateral Prefrontal Cortex | 45/46 | 60 | 4.32 | -57 | 21 | 24 |
| L Supplementary Motor Area | 6/8 | 142 | 4.23 | -21 | 12 | 57 |
| R Inferior Parietal Lobule | 40 | 319 | 5.76 | 54 | -66 | 36 |
| L Inferior Parietal Lobule | 40 | 281 | 5.51 | -48 | -78 | 33 |
| Study 2 (GLM 1b) |  |  |  |  |  |  |
| R Dorsal Anterior Cingulate | 6/8/9/32 | 936 | 5.27 | -3 | 35 | 46 |
| L Inferior Frontal Gyrus | 45/47 | 373 | 4.82 | -45 | 32 | -8 |
| R Inferior Frontal Gyrus | 47 | 268 | 5.06 | 39 | 23 | -11 |
| L Dorsolateral Prefrontal Cortex | 45 | 7† | 3.6 | -57 | 20 | 22 |
| L Middle Frontal Gyrus | 6/8 | 293 | 4.52 | -24 | 20 | 52 |
| L Posterior Cingulate Cortex | 31 | 100 | 5.12 | -6 | -40 | 34 |
| L Middle Temporal Gyrus | 21 | 38 | 4.07 | -60 | -31 | -8 |
| L Inferior Parietal Cortex | 39 | 285 | 5.57 | -39 | -70 | 40 |
| R Occipital Cortex |  | 120 | 4.9 | 42 | -73 | 34 |
| Study 3 (GLM 1c) |  |  |  |  |  |  |
| R Dorsal Anterior Cingulate | 6/8/9/32 | 472 | 5.28 | 9 | 23 | 40 |
| R Dorsolateral Prefrontal Cortex |  | 684 | 5.21 | 54 | 23 | 19 |
| Inferior Frontal Gyrus |  | * | 4.11 | 33 | 17 | -11 |
| L Dorsolateral Prefrontal Cortex |  | 671 | 5.23 | -51 | 20 | 19 |
| Inferior Frontal Gyrus | 47 | * | 4.43 | -33 | 26 | -5 |

*Note.* Regions are reported at a voxel-level of $P < .001$ , uncorrected and a whole-brain cluster corrected level of $P < .05$ , unless otherwise noted. \* Distinct peak within larger cluster. † Significant at $P < .05$ , small-volume corrected within a 10-mm spherical region of interest centered on the left dlPFC.

**Table S3.**
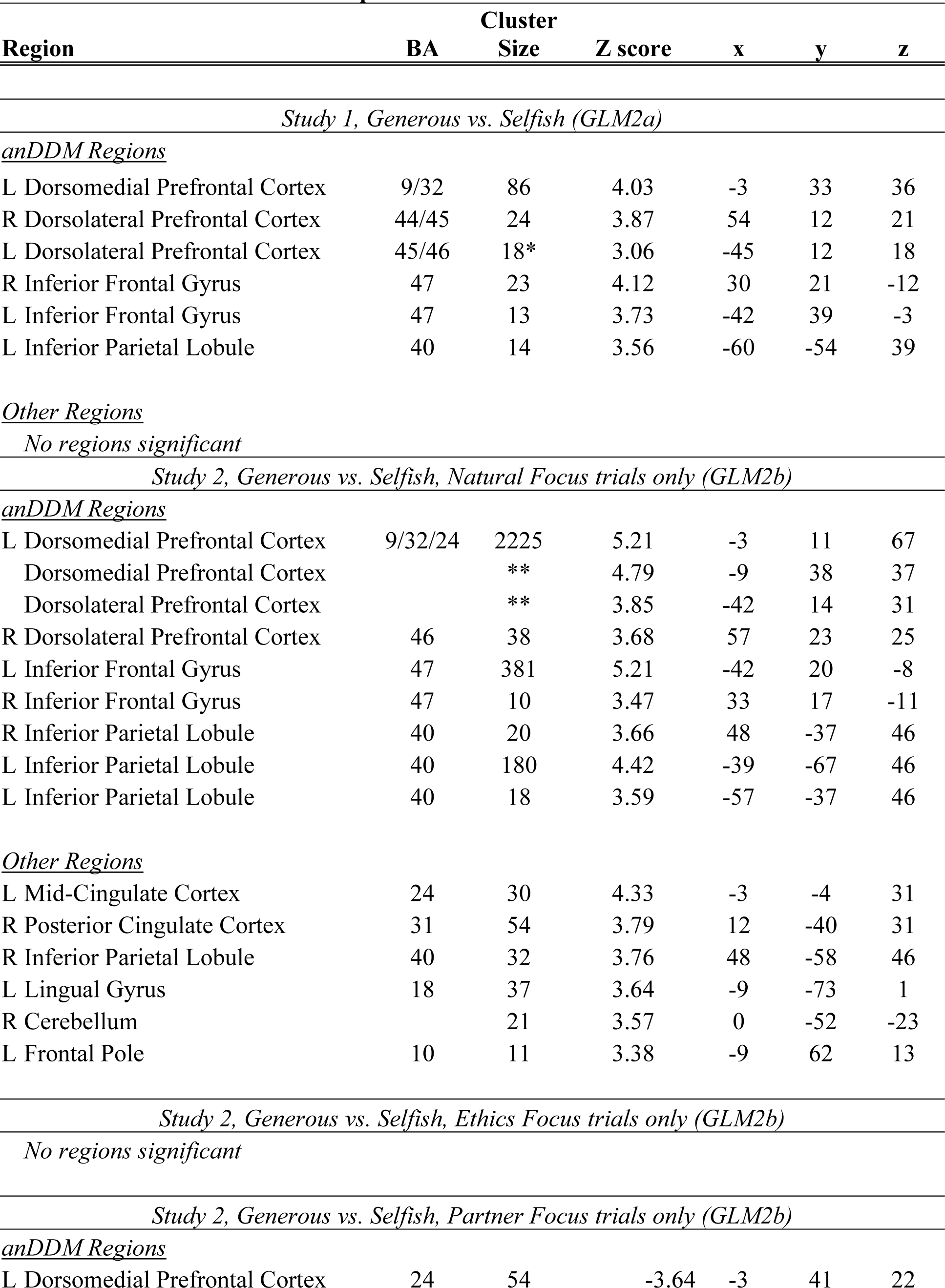
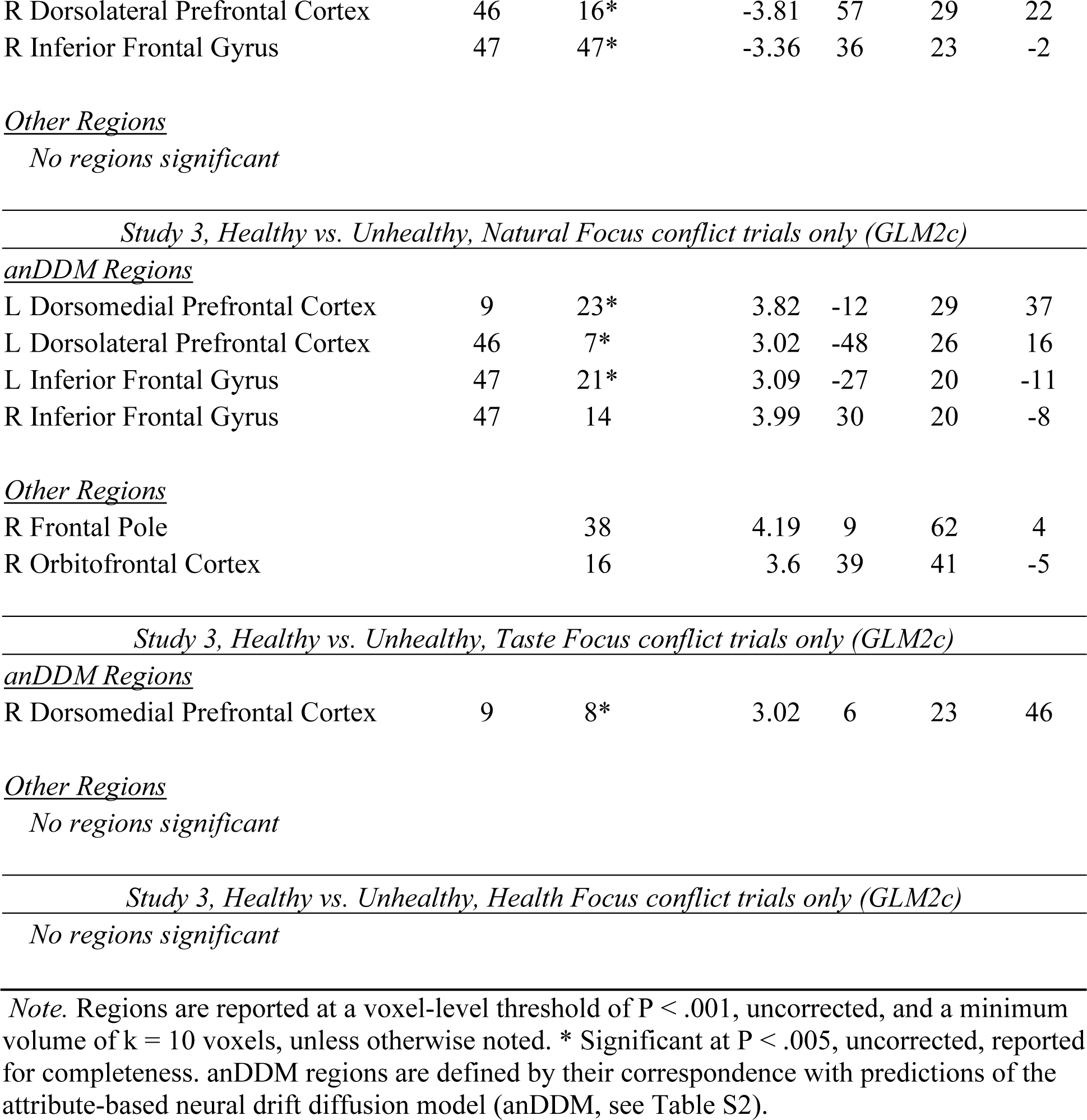
Differences in neural response for virtuous vs. hedonistic choices

| Table S5: Differences in neural response for virtuous vs. hedonistic choices |  |  |  |  |  |  |
| --- | --- | --- | --- | --- | --- | --- |
| Region | Cluster |  |  | x | y | z |
|  | BA | Size | Z score |  |  |  |
| *Study 1, Generous vs. Selfish (GLM2a)* |  |  |  |  |  |  |
| <u>anDDM Regions</u> |  |  |  |  |  |  |
| L Dorsomedial Prefrontal Cortex | 9/32 | 86 | 4.03 | -3 | 33 | 36 |
| R Dorsolateral Prefrontal Cortex | 44/45 | 24 | 3.87 | 54 | 12 | 21 |
| L Dorsolateral Prefrontal Cortex | 45/46 | 18* | 3.06 | -45 | 12 | 18 |
| R Inferior Frontal Gyrus | 47 | 23 | 4.12 | 30 | 21 | -12 |
| L Inferior Frontal Gyrus | 47 | 13 | 3.73 | -42 | 39 | -3 |
| L Inferior Parietal Lobule | 40 | 14 | 3.56 | -60 | -54 | 39 |
| <u>Other Regions</u> |  |  |  |  |  |  |
| *No regions significant* |  |  |  |  |  |  |
| *Study 2, Generous vs. Selfish, Natural Focus trials only (GLM2b)* |  |  |  |  |  |  |
| <u>anDDM Regions</u> |  |  |  |  |  |  |
| L Dorsomedial Prefrontal Cortex | 9/32/24 | 2225 | 5.21 | -3 | 11 | 67 |
| Dorsomedial Prefrontal Cortex |  | ** | 4.79 | -9 | 38 | 37 |
| Dorsolateral Prefrontal Cortex |  | ** | 3.85 | -42 | 14 | 31 |
| R Dorsolateral Prefrontal Cortex | 46 | 38 | 3.68 | 57 | 23 | 25 |
| L Inferior Frontal Gyrus | 47 | 381 | 5.21 | -42 | 20 | -8 |
| R Inferior Frontal Gyrus | 47 | 10 | 3.47 | 33 | 17 | -11 |
| R Inferior Parietal Lobule | 40 | 20 | 3.66 | 48 | -37 | 46 |
| L Inferior Parietal Lobule | 40 | 180 | 4.42 | -39 | -67 | 46 |
| L Inferior Parietal Lobule | 40 | 18 | 3.59 | -57 | -37 | 46 |
| <u>Other Regions</u> |  |  |  |  |  |  |
| L Mid-Cingulate Cortex | 24 | 30 | 4.33 | -3 | -4 | 31 |
| R Posterior Cingulate Cortex | 31 | 54 | 3.79 | 12 | -40 | 31 |
| R Inferior Parietal Lobule | 40 | 32 | 3.76 | 48 | -58 | 46 |
| L Lingual Gyrus | 18 | 37 | 3.64 | -9 | -73 | 1 |
| R Cerebellum |  | 21 | 3.57 | 0 | -52 | -23 |
| L Frontal Pole | 10 | 11 | 3.38 | -9 | 62 | 13 |
| *Study 2, Generous vs. Selfish, Ethics Focus trials only (GLM2b)* |  |  |  |  |  |  |
| *No regions significant* |  |  |  |  |  |  |
| *Study 2, Generous vs. Selfish, Partner Focus trials only (GLM2b)* |  |  |  |  |  |  |
| <u>anDDM Regions</u> |  |  |  |  |  |  |
| L Dorsomedial Prefrontal Cortex | 24 | 54 | -3.64 | -3 | 41 | 22 |
| R Dorsolateral Prefrontal Cortex | 46 | 16* | -3.81 | 57 | 29 | 22 |
| R Inferior Frontal Gyrus | 47 | 47* | -3.36 | 36 | 23 | -2 |

Other Regions *No regions significant*
| *Study 3, Healthy vs. Unhealthy, Natural Focus conflict trials only (GLM2c)* |  |  |  |  |  |  |
| --- | --- | --- | --- | --- | --- | --- |
| *anDDM Regions* |  |  |  |  |  |  |
| L Dorsomedial Prefrontal Cortex | 9 | 23* | 3.82 | -12 | 29 | 37 |
| L Dorsolateral Prefrontal Cortex | 46 | 7* | 3.02 | -48 | 26 | 16 |
| L Inferior Frontal Gyrus | 47 | 21* | 3.09 | -27 | 20 | -11 |
| R Inferior Frontal Gyrus | 47 | 14 | 3.99 | 30 | 20 | -8 |
| Other Regions |  |  |  |  |  |  |
| R Frontal Pole |  | 38 | 4.19 | 9 | 62 | 4 |
| R Orbitofrontal Cortex |  | 16 | 3.6 | 39 | 41 | -5 |

| *Study 3, Healthy vs. Unhealthy, Taste Focus conflict trials only (GLM2c)* |  |  |  |  |  |  |
| --- | --- | --- | --- | --- | --- | --- |
| *anDDM Regions* |  |  |  |  |  |  |
| R Dorsomedial Prefrontal Cortex | 9 | 8* | 3.02 | 6 | 23 | 46 |

Other Regions *No regions significant*
| *Study 3, Healthy vs. Unhealthy, Health Focus conflict trials only (GLM2c)* |
| --- |
| *No regions significant* |

*Note.* Regions are reported at a voxel-level threshold of $P < .001$ , uncorrected, and a minimum volume of $k = 10$ voxels, unless otherwise noted. \* Significant at $P < .005$ , uncorrected, reported for completeness. anDDM regions are defined by their correspondence with predictions of the attribute-based neural drift diffusion model (anDDM, see Table S2).

